# Charting the landscape of cytoskeletal diversity in microbial eukaryotes

**DOI:** 10.1101/2024.10.18.618984

**Authors:** Felix Mikus, Armando Rubio Ramos, Hiral Shah, Marine Olivetta, Susanne Borgers, Jonas Hellgoth, Clémence Saint-Donat, Margarida Araújo, Chandni Bhickta, Paulina Cherek, Jone Bilbao, Estibalitz Txurruka, Nikolaus Leisch, Yannick Schwab, Filip Husnik, Sergio Seoane, Ian Probert, Paul Guichard, Virginie Hamel, Gautam Dey, Omaya Dudin

## Abstract

Microbial eukaryotes are small and often resistant to standard labelling and imaging techniques, and therefore remain understudied – despite their critical ecological importance - with the exception of a few established models. Here, we use Ultrastructure Expansion Microscopy (U-ExM) to carry out high-resolution volumetric imaging of over 200 cultured planktonic eukaryotes across major lineages. By combining U-ExM with pan- and specific immuno-labelling, we reveal novel microtubule and centrin-containing elements and assign molecular identities to enigmatic cytoskeletal structures observed previously only by electron microscopy. Our investigation represents the first systematic survey of the extensive cytoskeletal diversity on display across the eukaryotic tree, including the major species groups of dinoflagellates, haptophytes, ciliates, euglenids, cryptomonads, and green algae. Our U-ExM approach extends to mixed environmental samples, paving the way for environmental cell biology at ultrastructural resolution and unprecedented scale, a crucial step towards understanding and protecting complex ecosystems in the face of biodiversity loss.

## Introduction

For over two centuries, the study of microbial eukaryotes (also called “plankton”; or “protists” for largely historical reasons, although these widely used terms remain imprecise and paraphyletic), has significantly expanded our understanding of the natural world. This voyage of microscopic exploration began in the late 17^th^ century with Antoine van Leeuwenhoek, who used one of the first microscopes to reveal a previously invisible world teeming with “animalcules”.^1^ It continued through the late 18^th^ and 19^th^ centuries, fuelled in large by the first global marine expeditions, most famously that of the HMS Challenger. ^2,3^ Inspired by samples taken by the Challenger during its 4-year-long expedition, Ernst Haeckel’s striking illustrations of thousands of species provided an initial framework for conceptualising eukaryotic diversity and evolution from common origins. ^4,5^ The early naturalists, including Haeckel and his contemporaries, driven by a desire to explore life’s diversity and uncover fundamental principles of cellular physiology, nonetheless lacked the tools to properly investigate cellular architecture.

Fast forward to the present day: low-cost sequencing and the ability to metagenomically profile environmental samples have revolutionised our capacity to catalogue genetic diversity across ecosystems, uncovering novel species and traversing the web of inter-species interactions at an unprecedented rate. ^6–8^ While these approaches have provided invaluable insights, they fall short of linking genotype with cellular architecture and phenotype, an essential aspect of understanding how eukaryotic cells function and interact in the context of the natural environment. Understanding cellular architecture goes beyond cataloguing compartments and cytoskeletal structures; it is necessary to understand how eukaryotic cells are organised, how they respond to environmental changes, and how they contribute to ecosystem dynamics in the face of global change. This knowledge gap is particularly critical for microbial eukaryotes, where most advancements in cell and molecular biology have been restricted to a few model organisms. Standard immunostaining protocols, which combine chemical fixation with antibody staining, often fail to uniformly label diverse organisms due to complex protective cell surface layers or walls and divergent physiology. Moreover, conventional light microscopes lack the resolution necessary to fully assess the subcellular complexity of eukaryotic cells, particularly those in the 5 to 50 μm size range that represent the bulk of microbial eukaryotic diversity. ^9,10^ Finally, although transmission electron microscopy (TEM), and more recently volume electron microscopy (vEM) of resin-embedded samples provide nanometre-scale resolution, the broad utility of these techniques is currently limited by throughput, cost, and the inability to directly label macromolecular structures of interest. ^11^

A promising solution to these challenges, with the potential to massively scale up microbial ultrastructure imaging, is expansion microscopy (ExM). ^12–15^ This recently developed method enables super-resolution imaging with conventional widefield and confocal microscopes through the isotropic expansion of hydrogel-embedded biological samples. ExM has found applications in animal models, cell lines, and tissues and its use across diverse eukaryotes is on the rise. ^16–20^ Parasitologists were early adopters, with striking results in systems ranging from Apicomplexa to trypanosomes. ^21–24^ ExM has even proven effective on species with mechanically robust cell walls of diverse composition, including Ichthyosporea, Corallochytrea, *Chlamydomonas*, *Giardia,* plant roots and pathogenic and non-pathogenic yeasts, often without the need for an extra cell wall digestion step. ExM is compatible with most standard chemical and cryogenic fixation protocols, allowing the protocol to be tailored to the needs of each specific system and biological question^25–31^.

Building on these advantages, ExM has the potential to overcome the limitations of current imaging methods and significantly scale microbial imaging efforts at a low cost. However, the adaptability and efficiency of ExM for ultrastructure imaging across a wide range of microbial eukaryotes remain largely unexplored. In this study, we applied the Ultrastructure Expansion Microscopy (U-ExM) protocol to over 200 aquatic microbial eukaryotic species, maximising diversity across the catalogues of two leading European culture collections; the Roscoff Culture Collection (RCC; https://roscoff-culture-collection.org/) and the Basque Microalgae Culture Collection (BMCC; www.ehu.es/bmcc). ^15,32^ Using a standardised set of dyes and antibodies targeting key cellular structures, we provide a glimpse into an unexpected and remarkable universe of cytoskeletal diversity, allowing us to discover novel structures and expand our understanding of the fundamental organising principles of eukaryotic cells. We further explore the potential of adapting U-ExM for use in complex environmental samples, laying the early groundwork for a better understanding of microbial interactions and communities in natural ecosystems.

## Results

### Volumetric imaging of ultrastructure across the eukaryotic tree

To thoroughly assess the efficiency and broad-spectrum compatibility of ExM for ultrastructure imaging of diverse microbial eukaryotes, we collaborated with the Roscoff Culture Collection (RCC) and the Basque Microalgae Culture Collection (BMCC), giving us access to a large set of aquatic species in near-axenic culture. We selected over 200 species covering the major eukaryotic groups and representing the diversity in the culture collections (Table S1). While cryofixation preserves cellular architecture with minimum artefacts, chemical fixation is quick, can be applied to larger sample volumes, is less resource intensive and requires less post-processing, allowing easy scaling to 200 species and application in field settings. To preserve overall cellular architecture, including cytoskeletal structures and membranous organelles, we opted for formaldehyde fixation of concentrated cultures, and in a few cases, cryo-preserved samples, on-site at the marine stations hosting the culture collections. The samples were subsequently analysed by U-ExM in our laboratories. Among the many ExM protocols developed over the last decade, Ultrastructural Expansion Microscopy (U-ExM) has been efficiently applied for cell biological analysis of a variety of microbial eukaryotes including apicomplexan parasites, fungi, and protists with varying physiology and cell surface biochemistry. ^23,25,28,29,31^ U-ExM is distinguished from most ExM protocols in using post-expansion immunolabelling, thereby enhancing epitope accessibility and improving staining efficiency. We applied U-ExM across the selected species (Figure S1A; Table S1), combining immunofluorescence of conserved cytoskeletal components with Hoechst 33342 or DAPI for DNA staining, and used protein pan-labelling NHS-ester dyes to evaluate overall cellular organisation. ^23,33^ This DNA and pan-labelling approach is exemplified using *Amphidinium carterae*, one of the most abundant benthic dinoflagellates found worldwide (Figure 1A). Further, we investigated conserved cytoskeletal structures across our species set, focusing primarily on microtubules (Figures S1B-1H).

**Figure 1:**
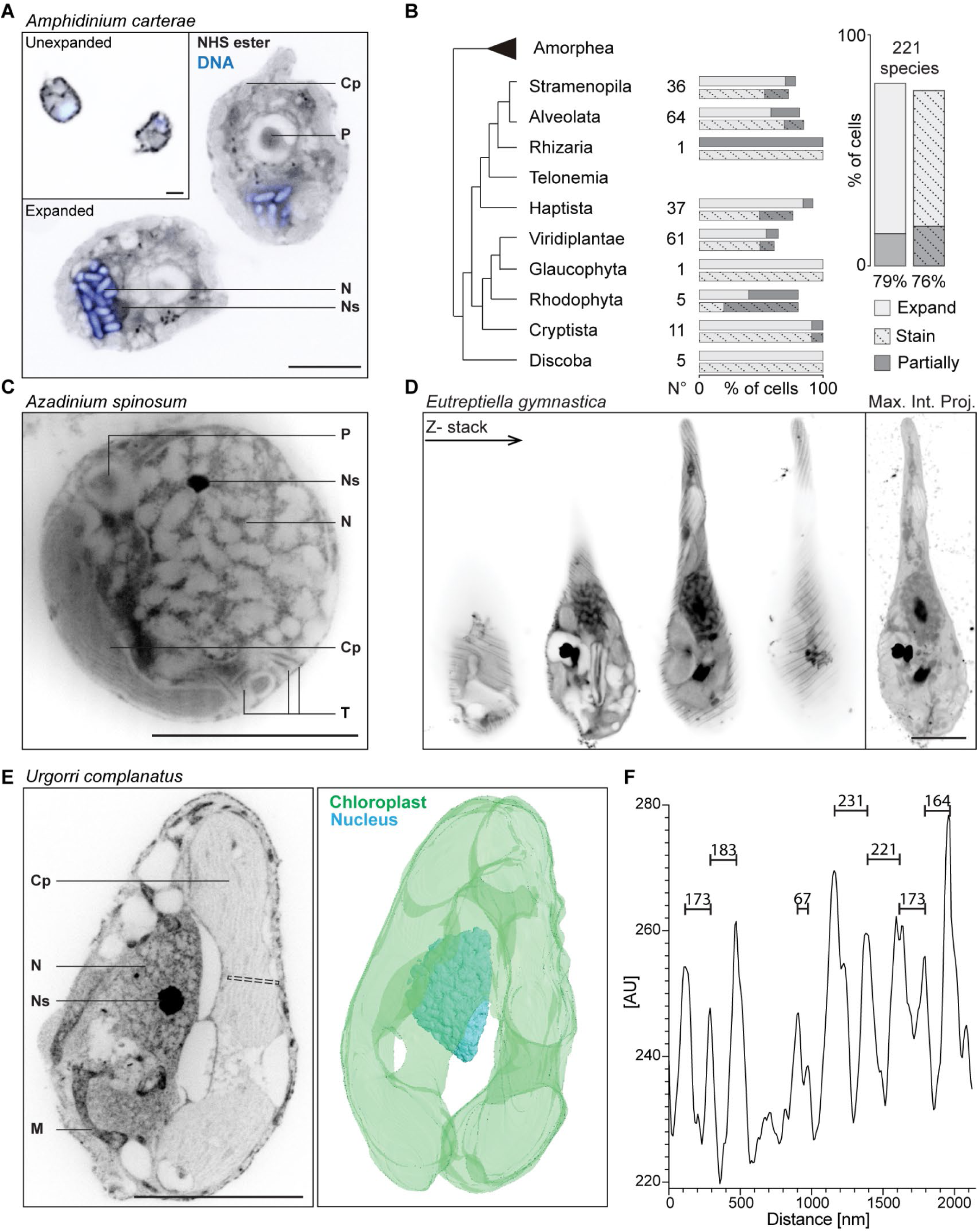
U-ExM as an efficient approach for imaging microbial eukaryotes. (A) Comparison of two median optical sections of an unexpanded and expanded dinoflagellate, *Amphidinium carterae* (not the same cell). Whole protein density is labelled with NHS-ester, DNA is shown in blue. (B) Summary of species processed for this study, sorted by clade. Light grey bars indicate successful expansion. Dark grey bars represent partial expansion (e.g., cell walls restricting parts of cells or individual cells expanding better than others), or instances where staining was present but incomplete. Striped background denotes the number of samples with staining. (C) Single optical section of *Azadinium spinosum* with the chloroplast (Cp), pyrenoid (P), nucleolus (Ns). The large gaps visible in the nucleus (N) are the chromosomes. Additionally, thin elongated rods close to the plasma membrane can be visualised, likely representing trichocysts (T). (D) Optical slices through a high-pressure frozen cell of *Eutreptiella gymnastica* and the corresponding maximum intensity projection. (E) In the cryptophyte *Urgorri complanatus,* the chloroplast (Cp), nucleus (N), and nucleolus (Ns) are easily recognised with NHS-ester staining. Furthermore, mitochondria (M) are localising in proximity to the furrow. Segmentation of the chloroplast and nucleus using NHS ester and Hoechst staining, respectively. (F) A line plot of the area indicated in E (dashed line) spanning over a part of the chloroplast. Peak to peak distances are indicated, representing thylakoid spacing. Scale bar, 5 µm. In the case of U-ExM images, the scale bar is adjusted for the expansion factor.

Out of the 221 species, 79% (175 species) successfully expand, and 77% (168 species) show at least partial staining with both pan-labelling and antibodies (Figures 1A and 1B; Table S1). To maximise the value of this work for the community, all raw images are available at (https://console.s3.embl.de/browser/culture-collections) and can be easily accessed using the MoBIE plugin in Fiji. ^34–36^ A curated selection of 60 species is also provided as a compiled single pdf file (Data S1; https://console.s3.embl.de/browser/culture-collections/data_s1%2F). Overall, no major lineage was completely refractory to the U-ExM protocol (Figure 1B). This suggests that the observed variability in expansion and staining efficiency is more likely due to species-specific physiological or structural properties, such as cell wall composition.

Using NHS-ester pan-labelling, we reveal the ultrastructure of various cellular organelles previously observed only with electron microscopy. For example, in the dinoflagellate *Azadinium spinosum,* known to have harmful effects on shellfish populations (Figure 1C), we can clearly distinguish the pyrenoid (P) and trichocysts (T). ^37,38^ Such ultrastructural details are not limited to two dimensions; we also obtain full volumetric information, as demonstrated for the euglenid *Eutreptiella gymnastica* (Figures 1D and S1B-S1E, Video S1), where the cell surface stripes, likely marking the pellicle, are visible at a variable interval below the optical resolution limit of conventional microscopes both with NHS-ester and tubulin staining (Figure S1C-S1E).

Further, chloroplasts identified by their thylakoid stacks, are discernible within the chloroplast of the cryptophyte *Urgorri complanatus* and can also be readily segmented (Figures 1E and S1H; Video S2). Using our approach, we find that the individual thylakoid stacks are spaced 180-240 nm apart (Figure 1F). Together, these results showcase U-ExM with NHS-ester pan-labelling, even in the absence of specific antibodies, as a versatile, scalable, and cost-effective tool to study subcellular organisation and organelle morphology across diverse plankton species in 3 dimensions.

### Diversity and complexity in the organisation of microtubule networks

Microtubule (MT) networks are ubiquitous across eukaryotes, playing crucial roles in regulating cellular architecture, motility, and division. Using anti-α and β-tubulin antibodies with broad species specificity, across our expanded sample collection, we observe significant diversity in MT organisation and structures among various species (Figure 2A). ^20,25,28^ Notably, the organisation of cortical MT arrays varies significantly between species as shown here in dinoflagellates and *Eutreptiella* species (Figure 2A). In many dinoflagellates, cells are divided into anterior epicone and posterior hypocone regions by the transverse girdle, or cingulum, which is delineated by two parallel MT bundles, often densely packed with MT arrays running perpendicular to the cingulum (Figure 2A). This MT arrangement contributes to dinoflagellate motility, particularly their characteristic “whirling” swimming motion, because the cingulum and sulcus serve as anchor points for flagellar attachment. ^39^ The epicone and hypocone MT arrays run obliquely or helically, converging at the cellular poles, with species-specific variability in the organisation of these regions. For instance, in *Gymnodinium impudicum* and *Protodinium* sp. both halves exhibit similar MT organisation with a somewhat similar spacing in cortical MT array (Figures 2B and 2C), whereas in *Takayama helix*, the epicone displays a distinct helical pattern (Figure 2A). ^40–42^ Distinct parallel cortical MT arrays were also observed in euglenid species *Eutreptiella gymnastica* (Figure 2A). These MT arrays likely underlie the pellicle, a complex structure composed of longitudinally arranged proteinaceous strips with MTs running beneath the plasma membrane providing structural support for morphological flexibility and adaptability to environmental conditions. ^43^ Moreover, in *E. gymnastica*, we identify a tubulin grommet surrounding the flagellar pocket collar (Figure S1F). ^44^ We identified two tubulin “spools” seemingly extended from a single MT in the dinoflagellate *T. helix* (Figures 2D and 2E; Video S3) at the nuclear periphery and at the posterior pole with a measured diameter of 3.67 μm and 2.85 μm, respectively (Figure 2F). We speculate that these spools may associate with the desmose described by Perret et al. in the dinoflagellate *Crypthecodinium cohnii*, potentially linking the basal body (kinetosome) and the centrosome in the context of cell division.^45^

**Figure 2:**
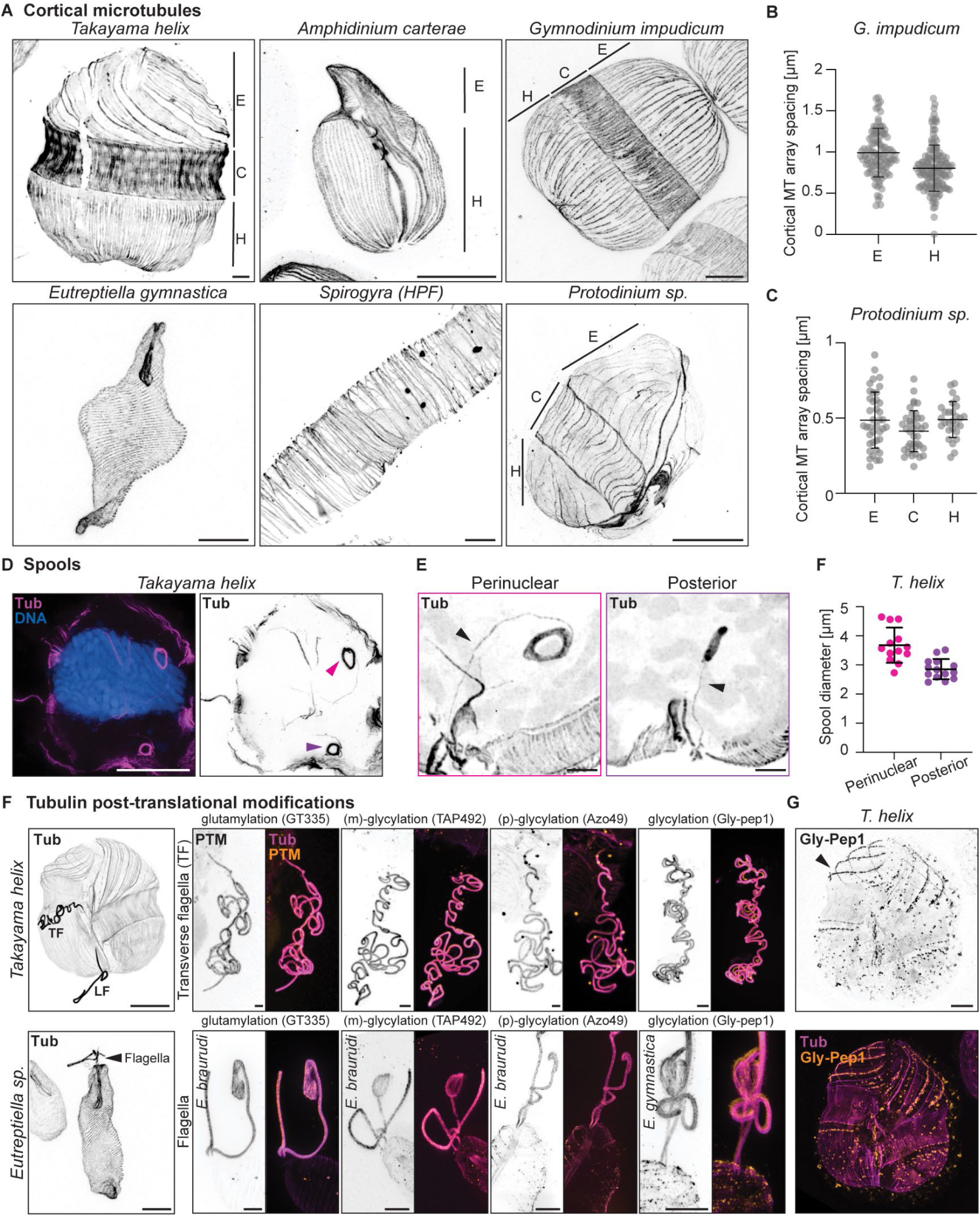
Diversity of microtubule organisation across eukaryotes. A) Maximum intensity projections showing cortical microtubule organisation in dinoflagellates, euglenoid, and *Spirogyra* species. In dinoflagellates, the epicone (E), cingulum (C) and hypocone (H) are indicated. B) Scatter plot showing MT array spacing in the epicone (E - 0.99 ± 0.29 μm, n = 57 microtubule array pairs) and hypocone (H - 0.80 ± 0.29 μm, n = 83 MT bundle pairs) in *Gymnodinium impudicum* (C), Scatter plot showing microtubule array spacing in *Protodinium* sp. (E – 0.48 ± 0.18 μm, C - 0.41 ± 0.13 μm H - 0.49 ± 0.11 μm, n = 40, 39 and 28 MT array pairs from the E, C and H regions respectively, collected from 3 cells each) (D) Maximum intensity projections showing microtubule spools (arrowheads, pink-perinuclear and magenta-posterior) in *Takayama helix*. (E), Perinuclear and posterior spool diameters are shown as a scatter plot (Perinuclear – 3.67 ± 0.6 μm, n = 13 spools, Posterior – 2.85 ± 0.33 μm, n = 14 spools). (F, G) Post-translational modifications of tubulin structures. Glutamylation and glycylation of dinoflagellate and euglenoid flagella (F) and glycylation of specific cortical microtubule bundles in *Takayama helix* (G). Dinoflagellate transverse flagellum (TF) and longitudinal flagellum (LF) are marked. All images are maximum-intensity projections of cells expanded and stained for tubulin (Tub), DNA (Hoechst). Scale bar 5 μm, adjusted for expansion factor. Error bars indicate mean ± SD (B, C and F).

Post-translational modifications (PTMs) of α/ß-tubulin, including glutamylation and glycylation, are important regulatory features of MT networks in model eukaryotes. To examine the conservation of this phenomenon, we stained both dinoflagellates (*T. helix*) and euglenids (*E. gymnastica* and *E. braarudii*), a group of eukaryotes in which PTMs have only been assessed in a recent study focused on the dinoflagellate *Ostreopsis* cf. *ovata*. ^46^ Using U-ExM, we find that the transversal flagellum of *T. helix* and the flagellum of *E. gymnastica* are modified by both glutamylation and mono- and poly-glycylation (Figure 2F).^47^ Notably, we reveal a distinct glycylation pattern in the cortical microtubules of the epicone and hypocone in *T. helix*, suggesting different functional roles within the cortex. This diversity of MT organisation across various dinoflagellate species opens up avenues for in-depth investigation of MT organisation as well as the relationship between MT array orientation and density and the arrangement of the protective amphiesma layer. More broadly, this analysis represents a step towards understanding the rules that govern the interplay between microtubule architecture and post-translational modifications across diverse eukaryotic taxa.

### Structural features of dinoflagellate chromosome and spindle architecture

Dinoflagellates possess unique chromosomal and spindle architectures, shaped by an unusual evolutionary history. ^48,49^ Unlike most eukaryotes, dinoflagellates exhibit permanently condensed, nearly crystalline chromosomes, a feature thought to be a consequence of the acquisition of viral and bacterial histone-like proteins, accompanied by a corresponding loss of their native complement of histones. ^50–52^ The distinctive banding patterns of dinoflagellate chromosomes, thus far only accessible to TEM, can now be effectively visualised using U-ExM – such as those in the dinoflagellates *Prorocentrum* and *Karenia*, 193 nm and 160 nm chromosomal band intervals, respectively (Figures 3A and 3B). The high throughput of our U-ExM imaging reveals cell-to-cell variability and species-specific differences in these banding patterns, which may reflect species-specific signatures or cell-cycle-dependent variations (Data S1) and merit deeper investigation.

**Figure 3:**
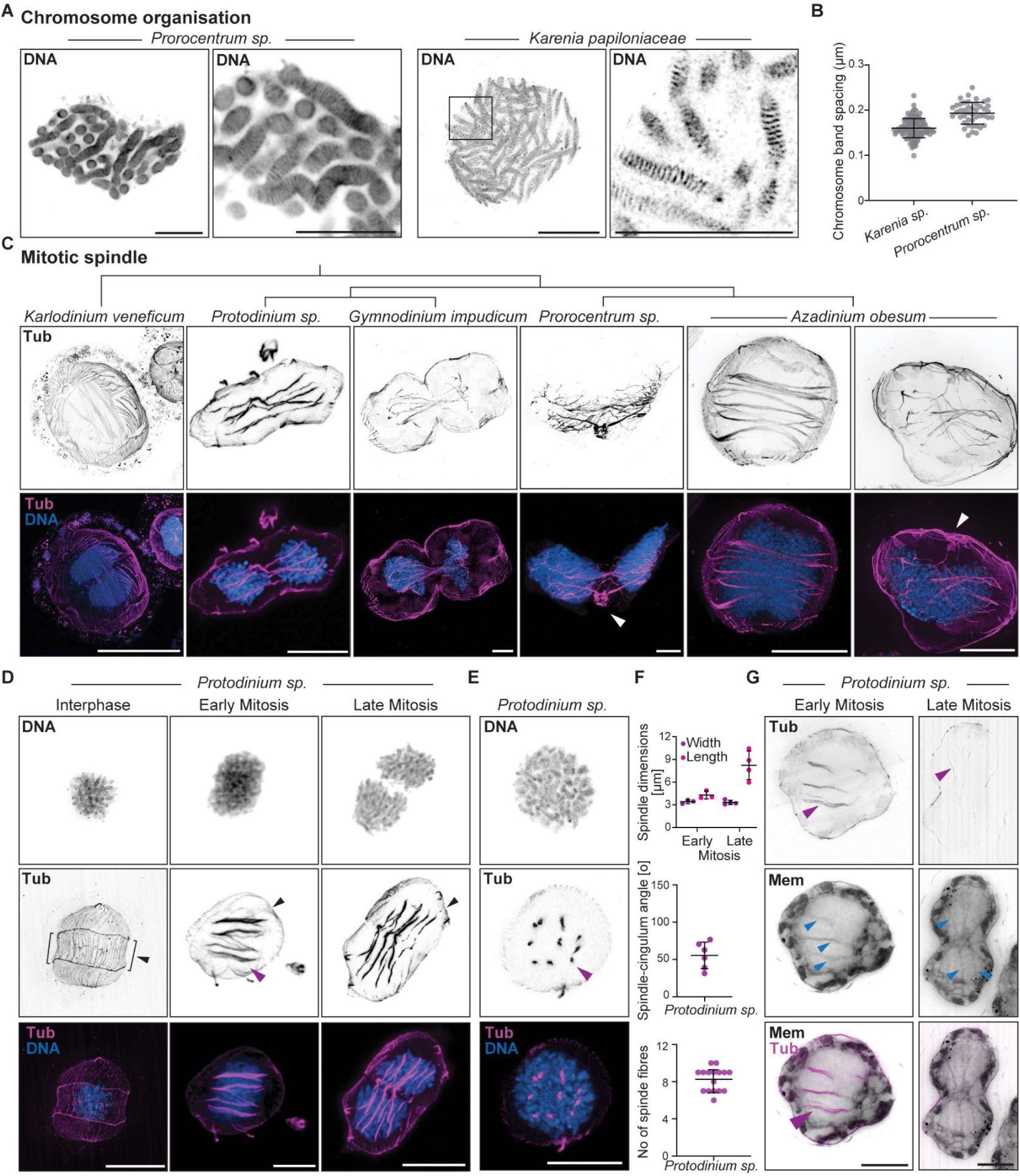
Chromosome and mitotic spindle organisation in dinoflagellates. A) Maximum intensity projections of dinoflagellate nuclei showing chromosome organisation (left) and anding pattern (right) for *Prorocentrum* sp. and *Karenia papilionaceae* (B) Scatter plot showing hromosome band spacing in *K. papilionaceae* (0.16 ± 0.021 μm, n = 116 chromosome band pairs from 2 uclei) and *Prorocentrum* species (0.193 ± 0.024 μm, n = 44 chromosome band pairs from 2 nuclei). C) Mitosis across different dinoflagellate species. White arrowheads indicate connections of the mitotic pindle to the flagellar apparatus. (D) Maximum intensity projections of *Protodinium* sp. cells in interphase nd mitosis (early and late) showing DNA and mitotic spindle (Tub) organisation, brackets and black arrows ndicate position and direction of cingulum and MT bundles (magenta arrowhead). (E) Cross section hrough a *Protodinium* mitotic spindle showing MT bundles (magenta arrowhead). (F) Spindle parameters, width (3.42 ± 0.24 μm, 3.325 ± 0.28 μm, n = 3 and 4 spindles respectively), height (4.29 ± 0.52 μm, 8.22 ± .89 μm, n = 3 and 4 spindles respectively), angle to cingulum (55.8 ± 17.8; n = 6 spindles) and number f spindle fibres (8.25 ± 1.25; n = 16 spindles) in early and late mitosis in *Protodinium* sp. (G) Maximum ntensity projections of *Protodinium* cells stained for membranes (Bodipy ceramide), showing intranuclear unnels (blue arrowheads) containing microtubule bundles (magenta arrowheads) in early and late mitosis. All images are maximum intensity projections of cells stained for tubulin (Tub), DNA (Hoechst) or membranes (Bodipy ceramide). Scale bar 5 μm, adjusted for ExM factor. Error bars indicate mean ± SD B and F).

This unique chromosomal architecture has important consequences for transcription, replication, and chromosome segregation during “dinomitosis”. ^53,54^ In this highly specialised form of closed mitosis, spindle microtubules form bundles that tunnel through the nucleus without penetrating the nuclear envelope (NE). The larger tunnels are crisscrossed by thinner tubules, forming a “nuclear net” first described using vEM in 2019. ^53^ How the spindle MTs attach to and segregate chromosomes without a direct interaction is not known, nor is the mechanism of nuclear division. Although our fixation strategy did not include a mitotic enrichment step, we fortuitously captured mitotic cells in various distantly related dinoflagellate species (Figure 3C).

Across these, we observe some variation in the number of spindle bundles and differences in their branching patterns and thickness (Figure 3C). In *Protodinium*, we find cells at various stages of mitosis (Figure 3D). The volumetric data allowed us to investigate spindle orientation and quantify the number of mitotic spindle fibres, which are visible in cross-sections (Figures 3D-3F). Additionally, U-ExM imaging provided pseudo-time series and quantitative measurements of spindle width and length, highlighting the potential for detailed studies of mitosis in cultured microbial eukaryotes even in the absence of synchronisation protocols (Figures 3E and 3F). Moreover, using the lipidic dye BODIPY-Ceramide, known to be compatible with U-ExM, we confirm the presence of the nuclear net, visible as membrane tunnels containing microtubules (Figure 3G). ^23,25,28^ We also observe pinching of the NE and intranuclear tubular network late in mitosis, just before cytokinesis (Figure 3G; Video S4), suggesting the involvement of contractile machinery, which we hypothesise could involve actin or other cytoskeletal elements. Together, these results could provide the initial basis for a thorough investigation of the evolutionary diversification of the mitotic machinery across dinoflagellates and their close relatives.

### Centrin as a cytoskeletal organiser in dinoflagellates

Next, we explored the complexity of cytoskeletal networks beyond microtubules, focussing on centrin—a key protein in centriole biology, known to localise at centrosomes, basal bodies of all flagellated or ciliated cell types - including mammalian cells, and several fibrous structures often associated with basal bodies in many flagellated or ciliated unicellular organisms. ^55–57^ Centrin is a well-conserved cytoskeletal protein found in all eukaryotic organisms and yet remains less understood in other cellular contexts. ^57–59^ Centrin belongs to the EF-hand superfamily of calcium-binding proteins, exhibiting 45–48% amino acid sequence identity with calmodulin. ^60^ Due to its ability to bind calcium, centrin is believed to play a role in forming or contributing to calcium-sensitive filamentous networks. ^61,62^ Additionally, centrin plays a crucial role in genome maintenance in animal cells, particularly through its involvement in the DNA repair process known as nucleotide excision repair(NER).^63^

In our investigation of dinoflagellates, we find that using the broad centrin antibody (20H5), centrin exhibits a variety of highly distinctive localisation patterns (Figure 4A). ^25,26,64,65^ For instance, we observe a centrin-based flagellar structure around which the transverse flagellum appears to coil, showcased here for *Protoceratium reticulatum* (Figure 4B, left panel). In certain dinoflagellates, the flagellum contracts upon exposure to increased intracellular Ca²⁺ and has been shown to consist of two fibres: the axoneme and the R-fibre.

**Figure 4:**
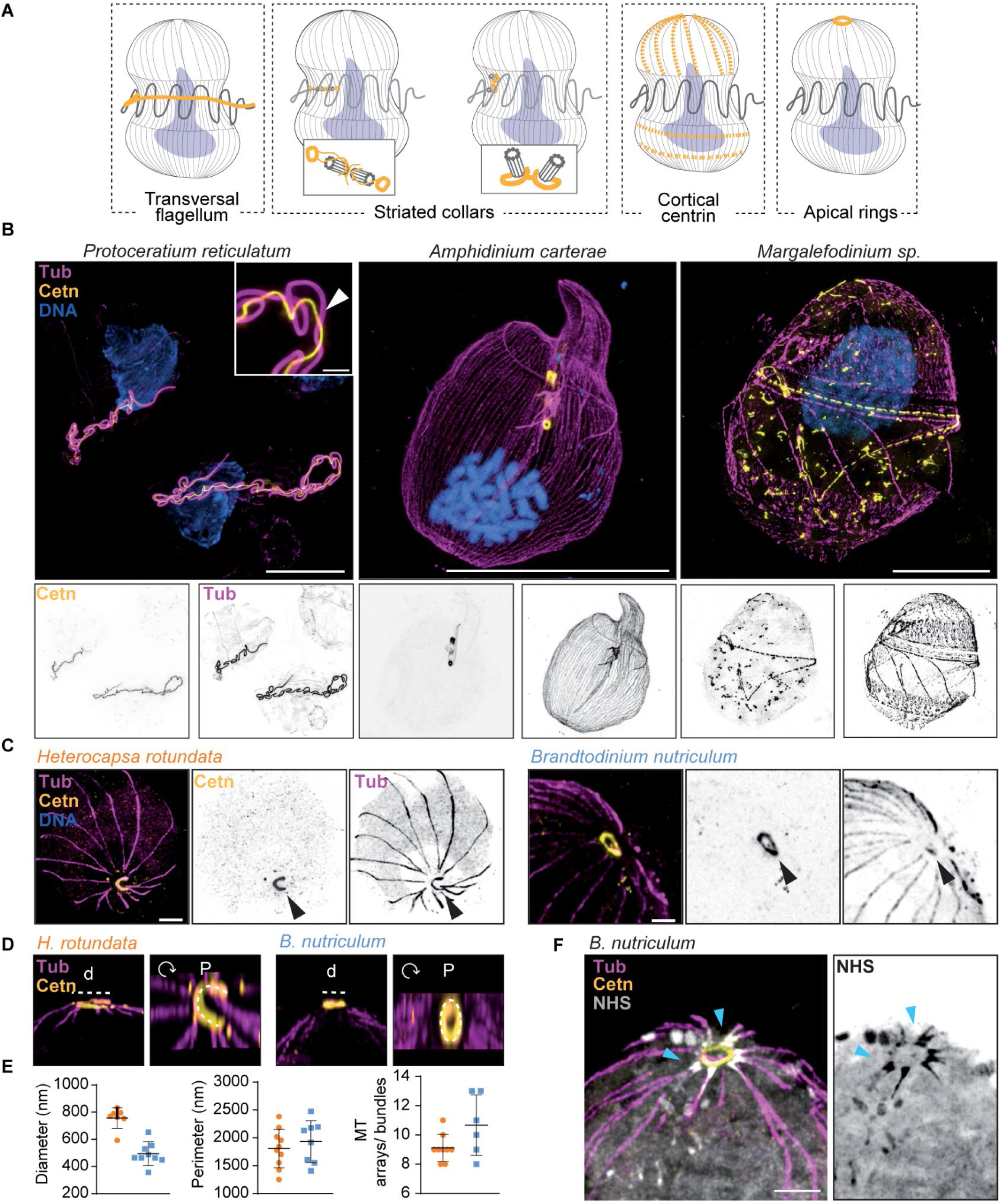
Dinoflagellates present high diverse centrin structures. (A) Scheme of the main centrin-containing structures found in dinoflagellates in this study. (B-C) Representative images of dinoflagellates *Protoceratium reticulatum*, *Amphidinium carterae* and *Margalefidinium* sp. immunostained with anti-tubulin (magenta) an anti-centrin (yellow) antibodies. In most of the species analysed, centrin showed a localization close-by the basal bodies, as seen in *Amphidinium carterae* (B). Examples of centrin being located below the plasma membrane forming a centrin mesh (B, middle), as part of the transversal flagellum (B, right) or forming an apical ring structure (C). (D) Zoom in on the centrin rings in *Heterocapsa rotundata* and *Brandtodinium nutriculum*. Graphs displaying the centrin ring diameter as well as its perimeter as indicated in the above cartoon. (E) Graph of the number of microtubule arrays (MT)/bundles emanating from the centrin ring (*H. rotundata*, 9±1 n = 9 cells; *B. nutriculum*, 11±2 n = 6 cells). (F) Representative image of an expanded *Brandtodinium nutriculum* cell mmunostained with anti-tubulin (magenta), anti-centrin (yellow) antibodies and NHS ester 594 (grey) highlighting densities connecting the microtubules bundles. Scale bars, 10 μm for (B), 1 μm for (C) and 5 μm for (F) adjusted to the expansion factor. Error bars indicate mean ± SD (D).

Our results suggest that the R-fibre, particularly in species exhibiting centrin-based flagella (Figure S2), either contains or co-localises with centrin, hinting towards centrin’s potential role in dynamic flagellar contractility and overall motility. ^66,67^ We also find that centrin localises in varying proximities to basal bodies depending on the dinoflagellate species, where it is anticipated to be part of the striated collar, as exemplified in *Amphidinium carterae* (Figure 4B, middle panel). The striated collar, surrounding the basal bodies, has been described in several dinoflagellates, where centrin fibres are proposed to anchor the flagella to the basal body complex, thus contributing to the organisation and stability of the flagellar apparatus.^68^ Additionally, we observe a distinctive cortical localisation of centrin in *Margalefidinium* sp. (Figure 4B, right panel). This cortical centrin pattern varies among the species we investigated (Figure S2; Data S1), suggesting it may confer species-specific structural integrity or motility adaptations and thus could serve as a tool for distinguishing between species or even different cell stages within a species. We also identify that centrin localises as apical rings in certain dinoflagellates (Figures 4A and S2). These apical centrin rings can be either open-sided, as seen in *Heterocapsa rotundata* (Figure 4C; Video S5), or closed, as seen in *Brandtodinium nutriculum* (Figure 4C; Video S6). Open-sided and closed rings are of varying diameter but tend to have a similar perimeter, averaging 1.8 µm (Figures 4D and 4E). When co-localising microtubules (MTs), centrin and the global proteome using NHS-ester pan-labelling, we observe that the centrin ring connects with an average of 9 or 11 MTs depending on the species (Figure 4E) via an as yet unidentified protein linker (Figure 4F, blue arrowheads), suggesting that the MT network is emerging from this uncharacterised structure.

Moreover, we find that the centrin ring in *Heterocapsa rotundata* colocalises with tubulin (Figure 4C, black arrowheads), while *Brandtodinium nutriculum* centrin rings show only a partial co-localization with tubulin (Figure 4C, black arrowheads). Within dinoflagellates, serial section TEM has previously shown the presence of a fibrous ring with emanating striated fibers right below the apical pore plate of the theca. ^68^ To our best knowledge, this description has been limited to two species, *Scrippsiella sweeneya* and *Heterocapsa pygmeae*, with the description of the ring as open sided in *Heterocapsa*, thus matching our observations in this study. These results are somewhat unexpected, as such apical open-sided/closed rings, composed of tubulin fibers, have been previously shown to be part of the apicomplexan conoid - a specialized, cone-shaped cytoskeletal structure found in some alveolates, comprising a diverse group of protists, including both dinoflagellates and apicomplexans. The molecular architecture and structure of the conoid have been extensively characterized in apicomplexan parasites, where a fully closed conoid tubulin-fibers-based structure exists, with centrin located in the pre-conoidal rings in *Toxoplasma* and absent in *Plasmodium.* ^69^ To our knowledge, this structure has never been described in dinoflagellates. ^70^ However, pseudo-conoids have been previously identified in colpodellids and perkinsids, which are sister clades of dinoflagellates, where they have developed an open conoid structure. ^70–73^ Although we cannot exclude the possibility that the apical centrin ring structures in dinoflagellates are unrelated to the known alveolate conoid, we anticipate that our results might have a significant impact on understanding the evolution of conoid structures in Alveolata and their role in the transition from a free-living to intracellular parasitic life cycle.^73,74^

### A centrin-based cytoskeleton is a defining feature of many microbial eukaryotes

Having identified the diverse cytoskeletal localisation and potential roles of centrin in dinoflagellates, we decided to extend our analysis to a broader range of microbial eukaryotes (Figure 5A; Table S1; Data S1). Once again, we observe centrin localising to distinct subcellular regions, with variations in its proximity to basal bodies, nuclear baskets, rootlet structures, and cortical networks (Figure 5A). For instance, in haptophytes such as *Dicrateria rotunda* and *Chrysotila roscoffensis*, centrin predominantly localises around the basal bodies (Figure 5B), with some centrin-positive arrays co-localising with tubulin, potentially indicating the presence of flagellar roots (arrowheads) (Figure 5B). In the halophilic green alga *Dunaliella salina*, we observe centrin forming a striking connection between the flagellar apparatus and the nucleus, suggesting a possible role in nuclear-cytoskeletal interactions (Figure 5C). These centrin fibers resemble the complex filamentous network described in *Chlamydomonas reinhardtii*, which extends from the nuclear surface to the proximal end of the flagella, known as the nucleus-basal body connector (NBBC). ^75^ This nuclear basket centrin localisation is not specific to *D. salina*, as similar patterns were also observed in other species, such as the raphidophyte *Chattonella subsalsa* (Figure 5C). Moreover, we also find centrin forming a distinctive cortical cytoskeletal network in cryptophytes as seen in *Rhodomonas* sp. and *Rhinomonas nottbecki* (Figures 5D-5F and S3A-S3B; Video S7). Although the centrin pattern superficially resembles that observed in ciliates, such as *Paramecium*, in which centrin plays a role in docking basal bodies to the plasma membrane, cryptophytes only have two cilia with no evidence of additional basal bodies at the cortex. The network’s arrangement and periodicity suggest a highly ordered structure, with a conserved periodicity between species as observed for both longitudinal (∼193 nm) and transversal axes (∼530 nm) (Figure 5E). In some species as *R. nottbecki*, centrin is observed forming hexagonal mesh/network (Figures 5F and S3) that can be quantified and probably indicates that centrin localises below the periplast, which is composed of hexagonal plates. ^76^ This centrin patterning shows some level of diversity among cryptophytes with *Urgorri complanatus* showing a full mesh with interspersed centrin rings, and *Cryptomonas curvata* missing any form of detectable peripheral centrin. Surprisingly, even between different isolates of the same species differences are notable since the afore-described network in *Rhodomonas* sp. was replaced by a more diffuse open-ringed pattern in a second culture (Figures S3A-S3B). Large amounts of centrin were also found along the length of the cryptophytes gullet and furrow (Figure S3C) where it might be involved in ejectosme release. Additionally, centrin localises as fibres with periodic striations at the flagellar rootlets, consistent with their involvement in contractile structures, as demonstrated in the green algae *Pyramimonas diskoicola* and *Microglena reginae* (Figure 5G, Video S8). Quantification of centrin striation periodicity (Figure 5H) revealed values around 200 nm for both species (Figure 5I), consistent with the 170-220 nm periodicity observed in system II fibers of *Tetraselmis subcordiformis*.^77^ Such centrin organization suggests evolutionary conservation in the structure and function of centrin-positive fibers.^78^ Taken together, our results reveal an extensive phylogenetic distribution of centrin-containing cytoskeletal fibres across microbial eukaryotes and suggest a novel structural cytoskeletal role for centrin in cryptophytes.

**Figure 5.**
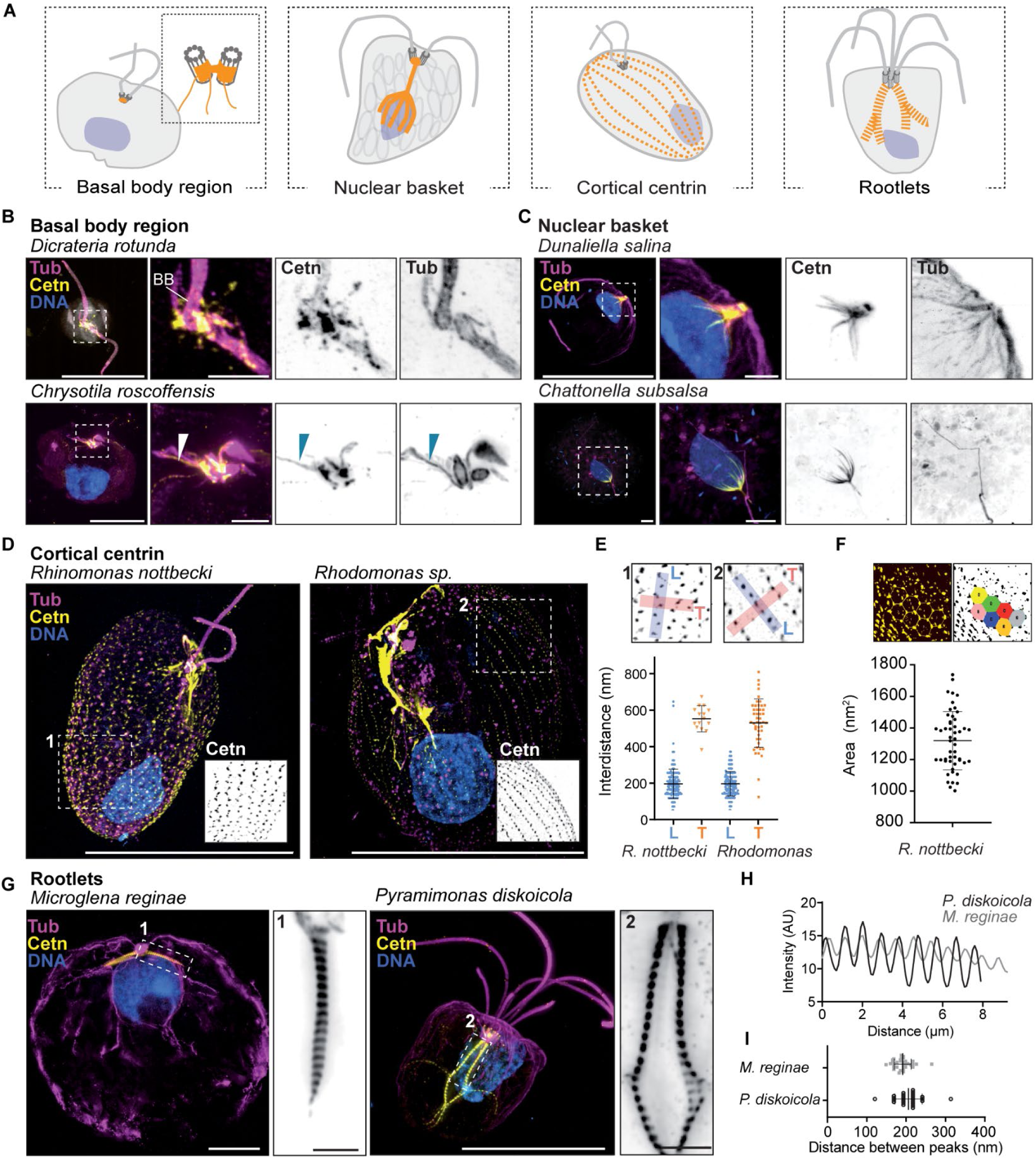
Centrin is the main component of different functional cytoskeleton structures. (A) Scheme showing the main centrin-structures found in this study. (B-H) Representative wield-field images of expanded samples immunostained with anti-tubulin (magenta) and anti-centrin (yellow) antibodies. (B) Centrin localises around the basal bodies (BB) in *Dicrateria rotunda* and *Chrysotila roscoffensis*. Centrin-positive arrays colocalising with tubulin that might correspond to flagellar roots are shown (arrowheads). (C) Centrin connects the flagellar apparatus with the nuclei in *Dunaliella salina* and *Chattonella subasalsa*. (D) Centrin forms a cytoskeleton network in some cryptophyceae. Representative images of *Rhodomonas* sp. and *Rhinomonas* sp. are shown. (E) Distance of the peak intensity of centrin along the longitudinal axis (L, in orange) or the transversal axis (T, in blue) in *Rhodomonas* sp. (L=192.6 ± 67; T=525.8±132, n = 7 cells) and *Rhinomonas nottbecki* (L=193 ± 81; T=550.1 ± 72, n = 5 cells). (F) Analysis of the area of the hexagons formed by the centrin network. (n = 4 cells; n=50 hexagons). (G) Images of the striated II fibres in *Pyramimonas diskoicola* and *Microglena reginae*. Note that in both cases the fibres are connected with the BB and maintain a periodic striation. (H) Plot profiles of the intensity of the centrin following the yellow signal shown in G. (I) Quantification of the distance between the intensity peaks of the centrin striation in *P. diskoicola* (orange; d=205.9 ± 35, n = 5 cells) and *M. reginae* (blue; d=191.3 ± 21, n = 6 cells). Scale bars = 10 μm, and 1μm for the insets, adjusted to the expansion factor. Error bars indicate mean ± SD (E, F, I).

### Cytoskeletal diversification and specialisation in mixed ciliate-dominated cultures

Many eukaryotes cannot be maintained under laboratory conditions, and axenic cultures fail to recapitulate the complexity of natural ecosystems that involve a complex web of inter-species interactions. As a first step towards ultrastructure imaging of complex environmental samples, we extended our U-ExM approach to mixed cultures derived from a sample collected in the shallow waters of Tokyo Bay (35° 37’ 53.1906’’, 139° 46’ 37.2072’’). We identify at least four ciliates—*Euplotes rariseta*, *Lacrymaria* sp., *Vorticella* sp., and possibly *Pseudourostyla*, along with one amoeba, *Flabellula* sp. (Figures 6A and S4). In *E. rariseta*, centrin localises at the basal bodies, with additional fibres intersecting between the para-oral membrane and the adoral zone of membranelles (AZM), highlighting a previously unreported centrin arrangement (Figures 6B and S5D). We identify tubulin bundles running orthogonally between peripheral dorsal microtubule strands, with an average spacing of 90 nm (Figures 6B, white lines, and 6C).

**Figure 6:**
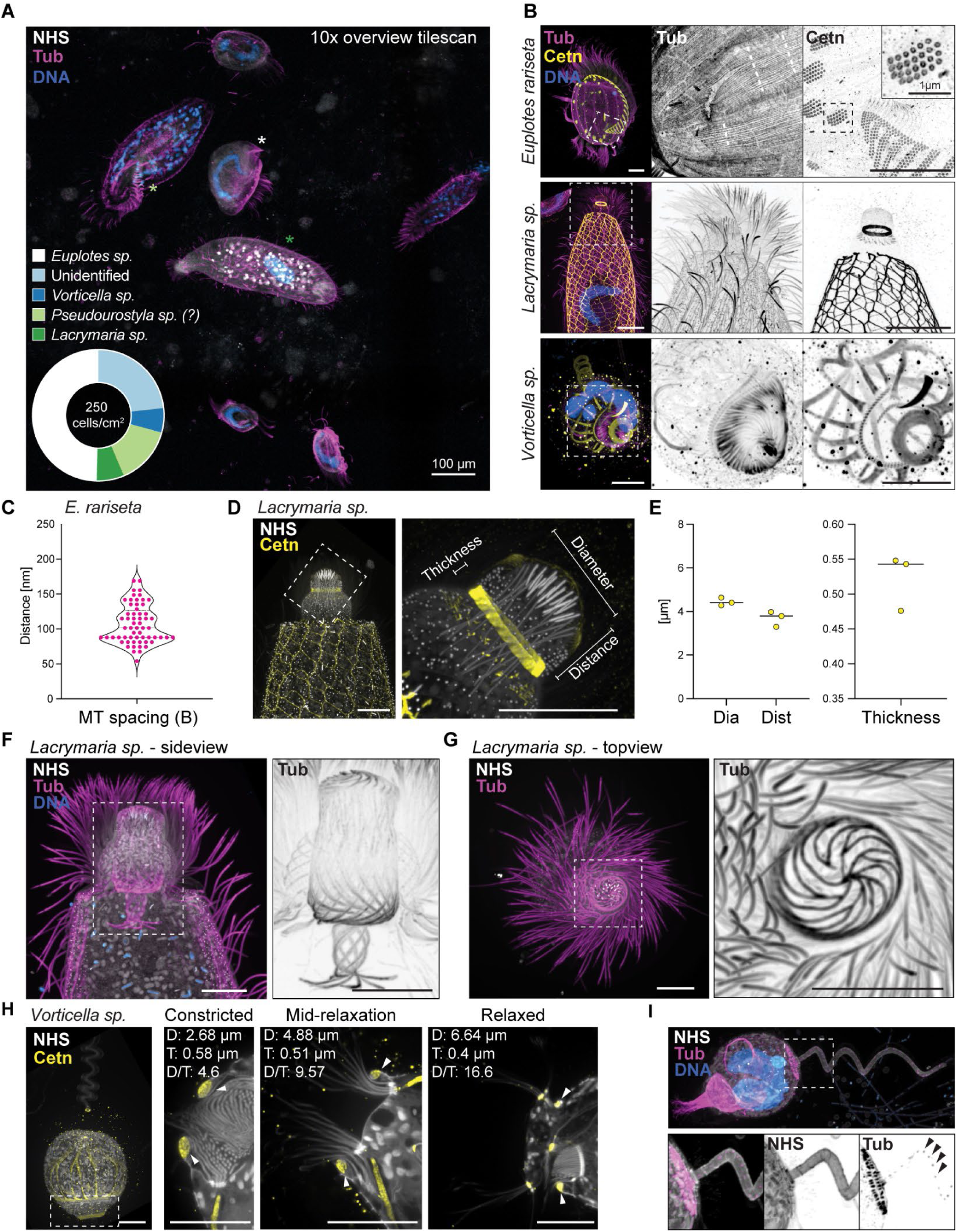
Expansion microscopy of ciliates collected from the environment. (A) Maximum intensity projection of an expanded sample imaged at low magnification reveals the species diversity and allows for the precise targeting of cells of interest. The pie chart displays the relative abundance of species in this particular gel. Cells of *Lacrymaria*, *Euplotes*, and *Pseudourostyla* are marked in the image with coloured asterisks. (B) Examples of *Euplotes rariseta, Lacrymaria* sp., *or Vorticella* sp. While *E. rariseta* exhibited the expected centrin localization to the numerous basal bodies of the cirri, bristles, and the adoral zone of membranelles (AZM), in both *Lacrymaria* and *Vorticella*, centrin forms intricate branched bundles and rings within their feeding apparatus. (C) Measurement of the distance between peripheral dorsal microtubule strands in *E. rariseta*, indicated by dashed white lines in (B). (D) The centrin ring localization in the mouth apparatus of *Lacrymaria*. (E) Measurements of the centrin ring in *Lacrymaria* indicating the distance from the cell tip (Dist), diameter of the ring (Dia), and thickness of the centrin signal (n = 3 cells). (F-G) The internal structure of tubulin bundles within the feeding apparatus of *Lacrymaria* sp. Resembling a vortex or chalice, cumulating in an aperture like at the cell tip. (H) The centrin ring in *Vorticella* changes in thickness (T) and diameter (D) dependent on the state of the cell, either contracted or with feeding cilia extended (relaxed). (I) Tubulin signals detected within the stalk of *Vorticella* as dots interspaced along the entire length. For all images, NHS-ester (grey), tubulin (magenta), centrin (yellow), DNA (blue). Scale bars, 100 µm (A) and 5 µm (B-H), adjusted for the expansion factor.

This arrangement supports the hypothesis that centrin and tubulin coordinate ciliary movement through complex interlinking networks. Imaging of various life stages of *E. rariseta* during vegetative growth, conjugation, and meiosis reveals large-scale macronucleus (MAC) changes, as well as, nuclear pore complexes organisation as pearls on a string localising between histone 3 enriched areas (Figures S5A and S5B). Notably, NHS ester staining highlights replication bands along the MAC (Figure S5A, magenta arrows), while NPC organisation remains constant, albeit depleted, in replication areas. Intranuclear bacteria were also detected in vegetatively growing cells during early culturing (Figure S6C), however, these bacteria were depleted from the culture before further detailed investigation could be performed.^79^ During meiosis, centrin staining revealed AZM remodelling, with rods replacing typical centrin rings.^80^

In *Lacrymaria*, centrin forms a peripheral fishnet-like network with prominent tubulin bundles spiralling between ciliary lines (Figure 6B). This is consistent with stainings previously shown for *Lacrymaria olor,* though the mesh size appears wider and more uniform throughout the entire body.^81,82^ We also observe a diagonally striated centrin ring, approximately 4 µm in diameter, located near the oral aperture (Figures 6D and 6E), probably supporting the structure of the cell’s feeding apparatus. The chalice-like vortex arrangement of tubulin bundles in the feeding structure is particularly striking, suggesting complex functional and structural roles during feeding (Figures 6F and 6G; Video S9). In *Vorticella*, we identify a prominent centrin ring at the edge of the oral apparatus, probably composed of several fibres, that increase in diameter during feeding (Figure 6H). Concurrently, we find that the thickness of this ring decreases, suggesting a sliding or stretching mechanism of centrin fibres (Figure 6H).

Surprisingly, we also detect tubulin-positive dots along the stalk at somewhat regular intervals, as well as a ring of tubulin dots at the stalk base (Figure 6I). The nature of these dots remains unclear; however, they may correspond to modified basal bodies as previously reported. These basal bodies consist of a circle of 9 single microtubules and are distributed along the length of the stalk, positioned adjacent to the alveolar-bearing side. ^83,84^ Further, centrin fibres extend from the oral apparatus towards the stalk most probably allowing for the contraction of the body (Figure 6H; Video S10). These structures, representing myonemes, have previously been shown to contain centrin by immuno-gold labelling.^85^ This study, however, presents the first 3D mapping of these structures in context. Overall, these findings demonstrate the straightforward application of U-ExM to complex cultures, opening the door to “environmental ultrastructure imaging” in the context of samples taken directly from the natural environment.

## Discussion

Despite the critical roles microbial eukaryotes play in Earth’s terrestrial and aquatic ecosystems, most cell biology research has focused on a few model organisms, leaving the vast diversity of this “invisible world” largely unexplored. This gap is partly due to technical limitations in visualising and studying the cellular architectures of microbial organisms. In this study, we demonstrate that Ultrastructure Expansion Microscopy (U-ExM) is a transformative tool for addressing this challenge, enabling high-resolution volumetric imaging of over 200 species of microbial eukaryotes across major lineages. We suggest that U-ExM could become a core cost-effective tool in protistology and microbial ecology, revealing previously unseen cellular structures and contributing to a more detailed understanding of how organisms function and interact within the context of their natural environments.

A key achievement of this work is the discovery and further cellular description of cytoskeletal structures across a broad range of species. Notably, we reveal tubulin-based networks and centrin structures, that exhibit remarkable species-specific variability. For instance, the identification of centrin apical rings, hinting at the presence of conoid-like structures in dinoflagellates, points to a potential evolutionary relationship between free-living protists and parasitic Apicomplexa, where the conoid has been extensively studied. The presence of this structure in non-parasitic species raises important questions about its evolutionary origins and functional diversification. A concern regarding the specificity of the anti-centrin 20H5 antibody remains, as the lack of genetic tools to deplete endogenous proteins limits direct validation. However, this antibody has been extensively used and characterised across a variety of microbial eukaryotes, including *Chlamydomonas*, *Giardia lamblia*, *Trypanosoma brucei*, *Leishmania major*, *Toxoplasma*, and *Tetrahymena*. ^23,25,86–88^ Notably, the antibody also recognizes human centrin. These findings can be attributed to the high sequence conservation of centrin, likely supporting the specificity of this antibody in our study. ^58,89^ Furthermore, our observations of spindle architecture in dinoflagellate mitosis, including variable nuclear envelope interactions and spindle organisation, provide new insights into the unique mitotic machinery of these organisms. The use of cultured species in this study paves the way for follow-up experiments, where many of the observed cellular structures can be further investigated with molecular tools to deepen our understanding of their evolutionary and functional significance. The scalability of U-ExM to diverse species, including those previously difficult to study, enables researchers to systematically analyse these structures and test specific hypotheses regarding their roles in microbial physiology, motility, and adaptation to environmental stress.

Our creation of an “atlas-scale” resource marks a significant advancement for protistologists and cell biologists alike. Over the course of 16 months, encompassing the fixation, expansion, and imaging of samples, we generated 1,670 image volumes from 221 species, resulting in approximately 4TB of data across ∼200 hours of microscopy. By providing a systematic, standardised dataset of ultrastructural images across a broad phylogenetic range, we aim to make this resource widely accessible to the scientific community. This resource serves as a reference for both experimental and computational studies, facilitating cross-species comparisons and enabling researchers to link cellular architecture with genetic and ecological data. Additionally, this work lays the groundwork for integrating U-ExM with other modalities, such as transcriptomics, proteomics, and metabolomics, offering a more comprehensive view of microbial eukaryotic biology ^10,90^. Future studies will benefit from incorporating life-cycle data to better understand how these structures evolve and change throughout organismal development.

Looking ahead, several areas will be critical for further expanding the utility and impact of U-ExM. First, the choice of dyes, antibodies, and fixation methods remains essential, as different species and cellular structures respond variably to these parameters. As is generally true for electron and light microscopy of preserved samples, the choice of fixation method in ExM is critical, with different macromolecules responding better to specific chemical fixatives. For example, aldehydes (formaldehyde or glutaraldehyde) stabilise proteins and a few lipids (amino lipids), while methanol is optimal for microtubules. ^91^ Some aspects of cellular architecture are better preserved through cryofixation. ^25^ The development of low-cost cryopreservation tools will also be crucial for increasing U-ExM’s accessibility to a broader range of laboratories, particularly those working with delicate environmental samples. Automation and scaling of both sample preparation and image acquisition will further facilitate high-throughput studies, reducing the time and effort required to process large numbers of samples. The application of AI and machine learning for image analysis represents another promising avenue, as automated segmentation and classification of cellular structures could allow researchers to rapidly process large datasets, making it easier to detect novel structures or identify species-specific patterns. Such approaches also enable real-time monitoring of complex environmental samples, where microbial communities exhibit high diversity and dynamic interactions. When integrated with advanced image analysis, U-ExM holds significant potential for environmental monitoring, particularly in assessing ecosystem health and biodiversity in the context of climate change.

In conclusion, U-ExM emerges as a powerful, cost-effective, versatile, and scalable method for imaging the ultrastructure of microbial eukaryotes in 3D. Its capacity to reveal fine cellular details at high throughput provides an essential tool for advancing our understanding of microbial diversity and the molecular mechanisms underlying their cellular organisation. As we continue to refine and apply the protocol, we expect U-ExM to play a pivotal role in future discoveries across cell biology, protistology, and microbial ecology, generating new hypotheses about the evolution and functional diversity of microbial life on Earth.

## Acknowledgements

We would like to thank all members of the Dey, Schwab, Centriole and Dudin labs, as well as the EMBL and external members of the TREC consortium, for many discussions and sustained feedback. We would like to thank the Roscoff Marine Station and the Plentzia Marine Station, University of the Basque Country (PiE-UPV/EHU), for hosting and supporting our work. We are grateful to Elisabeth Hehenberger and Buzz Baum for their critical reading of the manuscript, and to Carsten Janke for generously providing the antibodies targeting tubulin PTMs. Special thanks to Dirk Scholz for assistance with the Nikon NSPARC system and Niccolò Banterle for help with STED microscopy. We further acknowledge the support of the EMBL Mobile Lab Facility and the TREC expedition, as well as the Electron Microscopy Core Facility (EMCF) and the Advanced Light Microscopy Facility (ALMF). We further thank Pierre Gönczy and the Gönczy lab for fruitful discussions.

## Author contributions

Conceptualisation: OD, GD

<u>Methodology:</u> FM, ARR, HS, MO, SB, JH, CSD, MA, CB, PC, NL, YS, FS, SS, IP, PG, VH, GD, OD

<u>Software:</u> JH

<u>Validation:</u> FM, ARR, HS, MO, SB, JH, CSD, MA, CB, PC, NL, YS, FS, SS, IP, PG, VH, GD, OD

<u>Formal analysis:</u> FM, ARR, HS, PG, VH, GD, OD

<u>Investigation:</u> FM, ARR, HS, MO, SB, JH, CSD, MA, CB, PC, NL, YS, FS, SS, IP, PG, VH, GD, OD

<u>Resources:</u> FM, ARR, HS, MO, SB, JH, CSD, MA, CB, PC, JB, ET, NL, YS, FS, SS, IP, PG, VH, GD, OD

<u>Data Curation:</u> FM, ARR, HS, PG, VH, GD, OD

<u>Writing - Original Draft:</u> FM, ARR, HS, PG, VH, GD, OD

<u>Writing - Review & Editing:</u> FM, ARR, HS, MO, SB, JH, CSD, MA, CB, PC, JB, ET, NL, YS, FS, SS, IP, PG, VH, GD, OD

<u>Visualisation:</u> FM, ARR, HS, PG, VH, GD, OD

<u>Supervision:</u> PG, VH, GD, OD

<u>Project administration:</u> GD, OD

<u>Funding acquisition:</u> PG, VH, GD, OD

## Data availability statement

All microscopy datasets, which represent the resource provided by our study, are publicly hosted and openly accessible at S3 storage bucket: ^29–31^. This U-ExM images resource is also accessible as a MoBIE project at: https://console.s3.embl.de/browser/culture-collections. A curated list of images of 60 different species (Data S1) can also be found as a single pdf file at: https://console.s3.embl.de/browser/culture-collections/data_s1%2F.

## Code availability statement

All code necessary to upload the data as a MoBIE project can be found alongside the totality of the metadata in the following GitHUB repository: https://github.com/Dey-Lab-Team/culture-collections.

## Supplementary Figures

**Figure S1:**
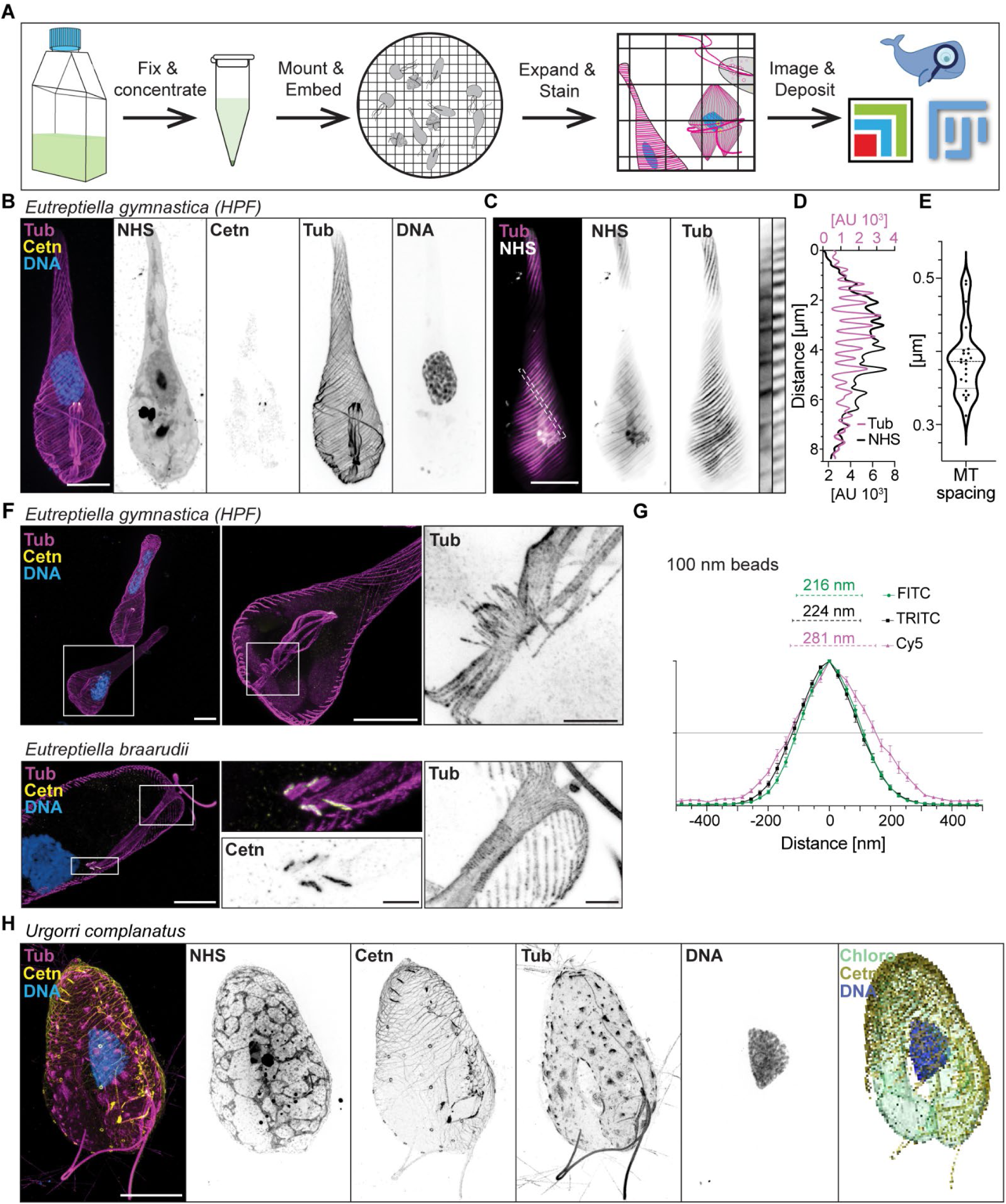
A streamlined U-ExM approach allows for fast high-resolution 3D acquisition. (A) Overview of the workflow starting with fixation of cultures, followed by concentration and by centrifugation before mounting onto coverslips and embedding into the expansion gels. After expansion, samples were stained with a standardised antibody and dye set, imaged and all images uploaded to the FIJI-based MoBIE plugin. (B) Maximum intensity projection of *Eutreptiella gymnastica* as in Fig. 1D with tubulin, centrin, and DNA staining shown. (C) Narrow maximum intensity projection representing 1 µm depth of Fig. 1D and Supp. Fig. 1B. The dashed line indicates areas cropped and shown for NHS ester and tubulin on the right edge. (D) Measurements of tubulin and NHS ester signals in 4 cells in two separate areas each. (E) Distances between tubulin peaks measured in D with an average spacing of 380 nm. (F) Comparison of the flagellar pocket of two *Eutreptiella* species. Cropped areas show orthogonal microtubules at the collar in both species. (G) Averaged intensities of eight 100 nm beads were measured in the three indicated channels. Distances at half-maximal intensity are noted. Note, that here no expansion factor needed to be factored in meaning that measurements down to 70 nm would be well within the resolution limit when expanding samples four-fold. (H) *Urgorri complanatus* from Fig. 1E, shown with staining for tubulin, centrin, and DNA. Segmentation of centrin signals, the DNA signal and chloroplast are also displayed.

**Figure S2:**
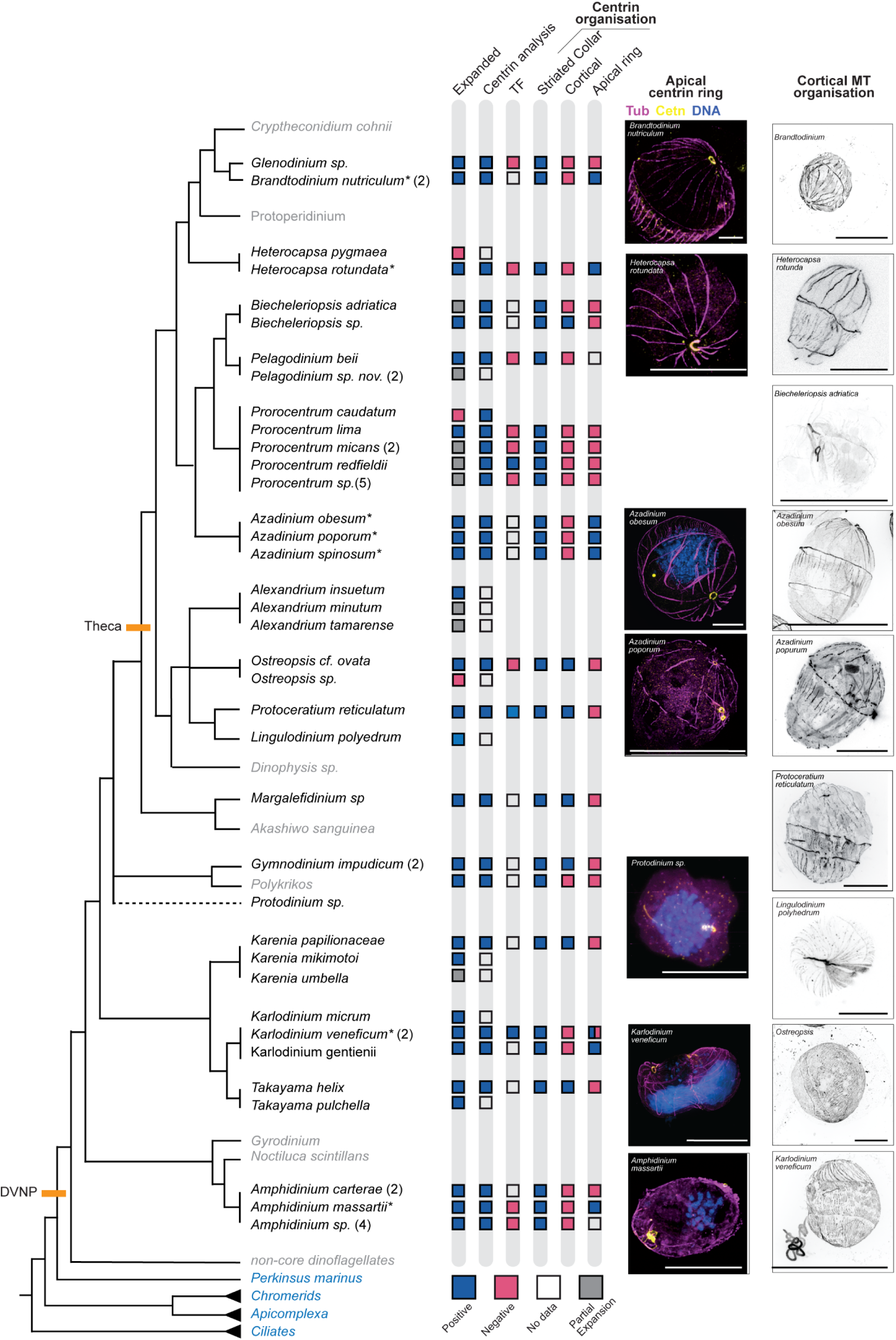
Cytoskeletal diversity across dinoflagellates. The cladogram shows the phylogenetic relationships of analysed dinoflagellate species based on previous literature. ^48,92,93^ Species included in the analysis are labelled in black, and others are marked in grey. Other alveolate relatives, Apicomplexa and ciliates, are marked in blue. The position of *Protodinium* sp. within the core dinoflagellates remains unresolved. Key transitions in dinoflagellate evolution are marked on the tree in orange. The squares on the right show profiles of centrin and tubulin organisation across the analysed set, positive (blue), partial expansion (grey), negative (pink) and no data (white). Maximum intensity projections highlight a few centrin and tubulin organisations in selected species. Scale bar, 5 μm.

**Figure S3:**
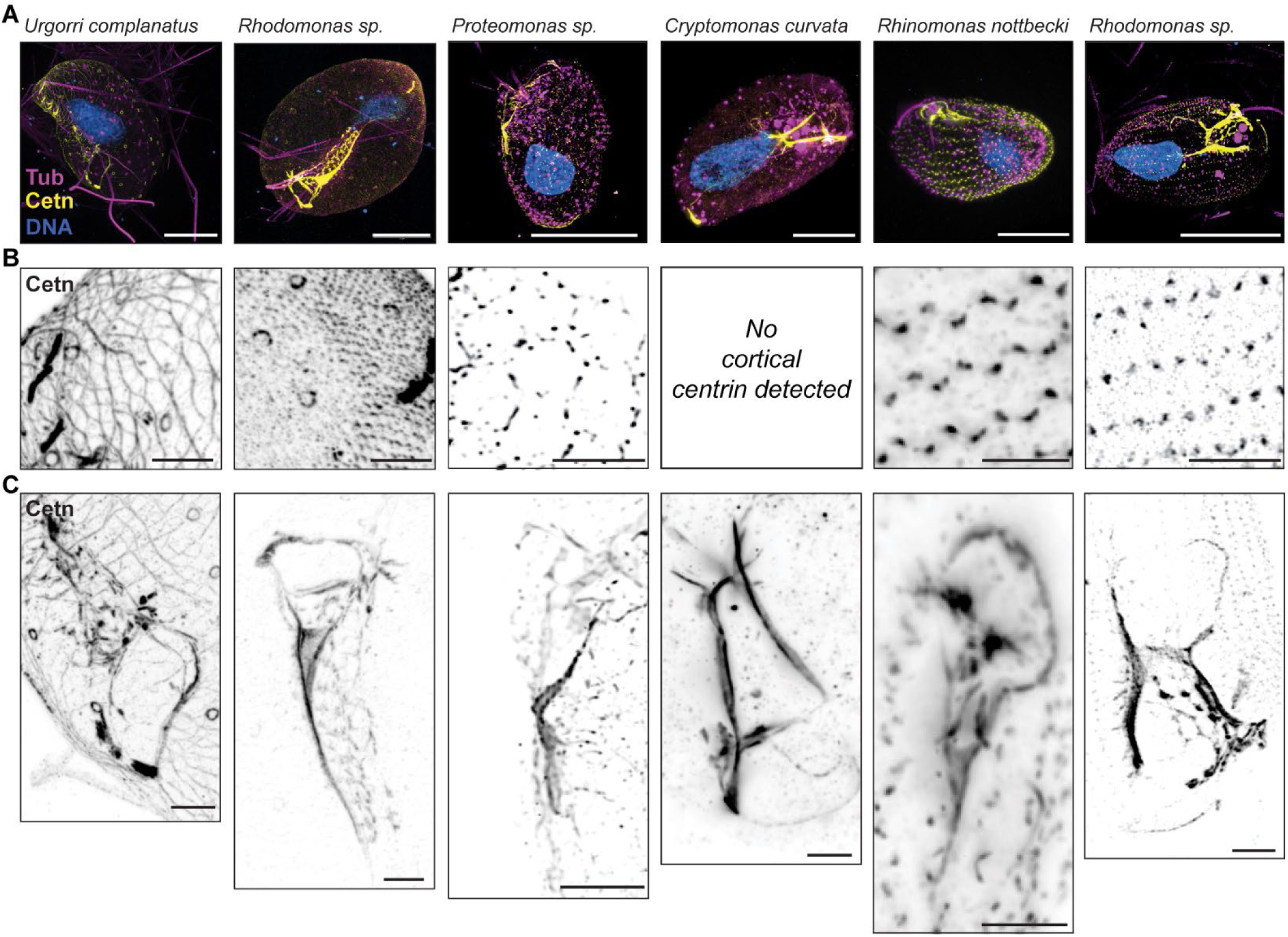
Cortical centrin mesh arrangement in cryptophytes. (A) Maximum intensity projections of cryptophyte *Urgorri complanatus, Proteomonas, Cryptomonas curvata, Rhinomonas nottbecki*. And two *Rhodomonas* species stained for tubulin, centrin and DNA. (B) Zooms into the cortical centrin network and (C) into the furrow. Scale bars 5 μm, adjusted for the expansion factor.

**Figure S4:**
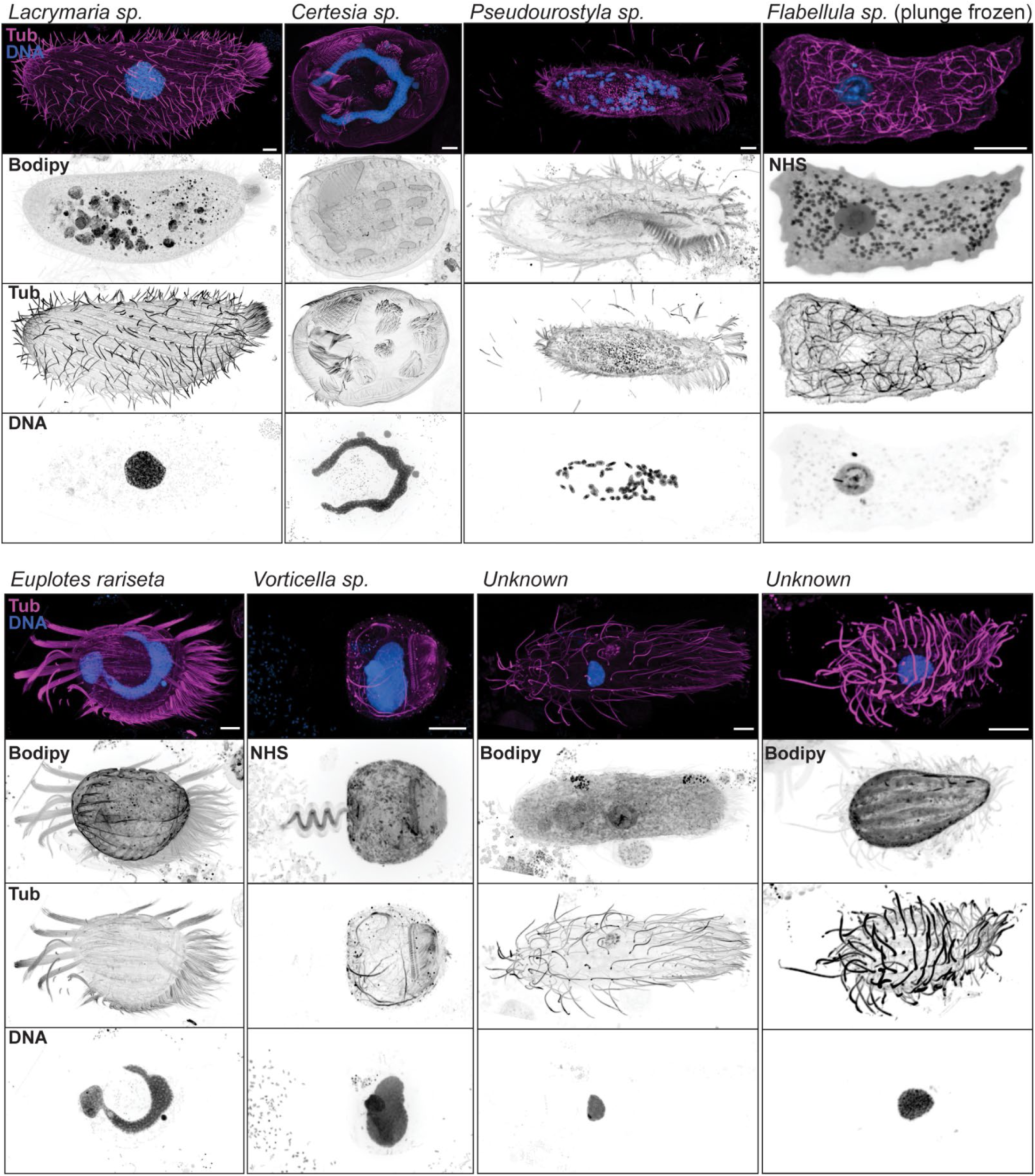
Visualising eukaryotic diversity from an environmental sample. Examples of cells of distinct species imaged within the environmental sample collected in Tokyo Bay. Only *Flabellula* sp. and *Euplotes rariseta* were confirmed by 18S sequencing. Other species were attempted to be identified by morphology. Note that in *Pseudourostyla* sp. and, in some cases, in *Vorticella* sp. expansion was incomplete resulting in some artefacts. All scale bars indicate 5 µm adjusted for the expansion factor. In all merge images, tubulin is displayed in magenta, centrin in yellow, DNA in blue.

**Figure S5:**
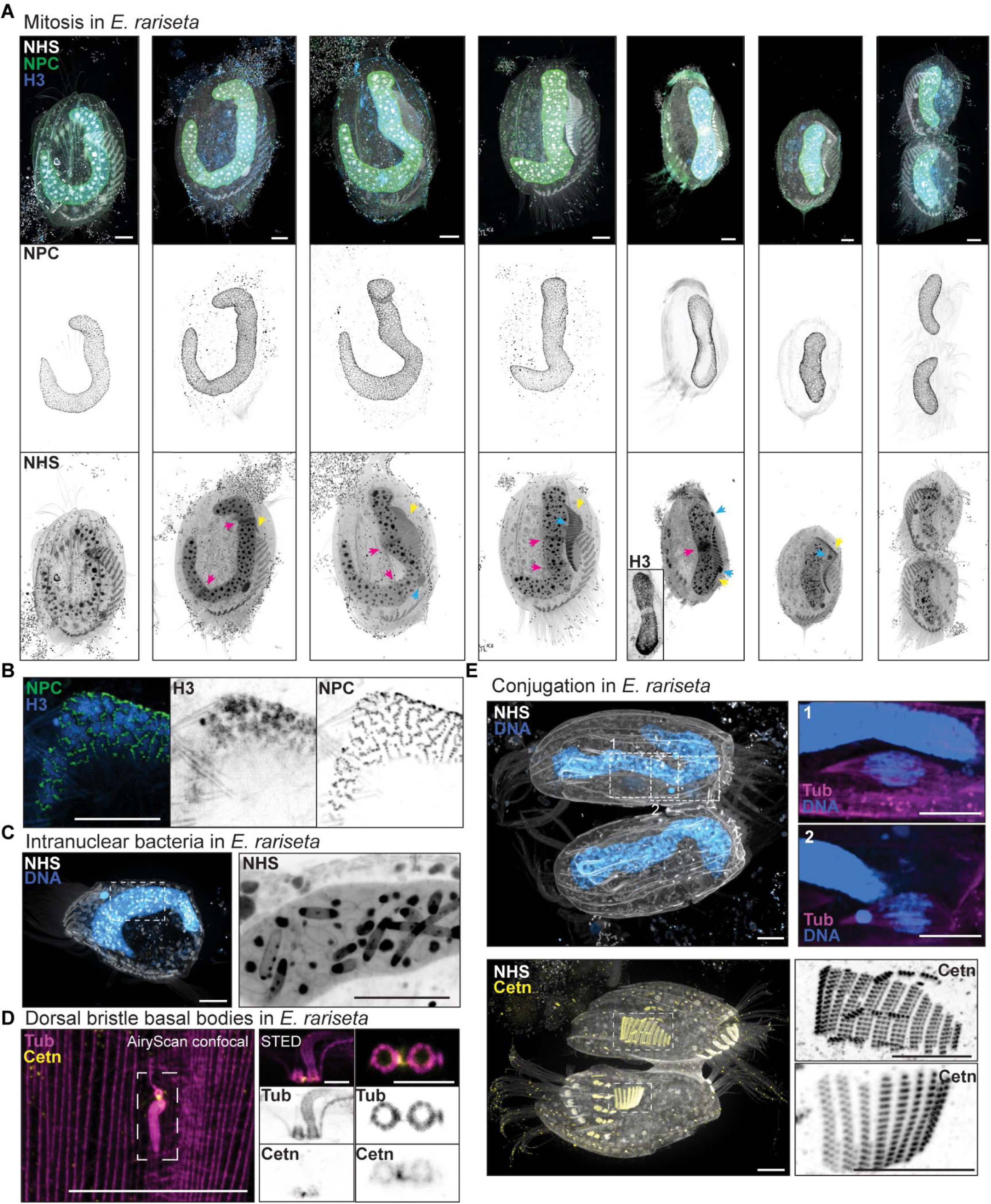
Mitosis in *Euplotes* sp. (A) Pseudo timeline of the mitotic life cycle of *E. rariseta*. Nuclear pore complexes (NPCs) were labelled with the nucleoporin antibody mab414 (which does not recognise NPCs within the micronucleus; MIC) indicating the nuclear envelope. Cells were staged by the presence or absence of the replication bands (magenta arrows) in NHS ester staining as well as by the state of AZM replication (yellow arrow) and MIC division state (blue arrow). An inset is shown for Histone H3 staining showing the depletion of H3 at the replication band while NPC distribution appears unaffected. Maximum intensity projections are shown. (B) A close-up of a macronucleus (MAC) in interphase is indicated in A. For H3, a sum intensity projection is displayed indicating regions of enrichment. For NPCs, a maximum intensity projection indicates their presence only between H3-rich areas as densely spaced rows reminiscent of Turing patterns. (C) During the early stages of culturing, *E. rariseta* often showed intra-MAC structures suspected to be bacteria. (D) STED microscopy of tubulin and centrin at dorsal bristle basal bodies. An overview, taken at an AiryScan confocal microscope, shows their localisation between parallel arrays of dorsal microtubule bundles. Lateral and frontal views taken at the STED show the nine-fold symmetry of the basal body as well as the left transverse (LTR) and right radial ribbon (RRR) or post-cilliary ribbon in tubulin staining as well as a centrin connector linking the two barrels at the lower end with a biased localisation to the basal body not supporting a cilium (500 nm scale bar). (E) Two examples of conjugation events in *E. rariseta*. Dividing MIC can be seen, while rod-like, densely spaced centrin signals clearly define the reformation of the AZM centrioles. If not mentioned otherwise, scale bars indicate 5 µm adjusted for the expansion factor. In all merge images, tubulin is displayed in magenta, centrin in yellow, and DNA in blue.

## Video Legends

**Video S1:** *Eutreptiella gymnastica* stained with NHS-ester staining (grey) and DNA (blue). Scale bars are not adjusted to the expansion factor.

**Video S2:** *Urgorri complanatus* stained with NHS-ester staining (grey) and DNA (blue). Scale bars are not adjusted to the expansion factor.

**Video S3:** *Takayama Helix* cell showing the tubulin spools (magenta) localised in the posterior and the perinuclear area and the cortical pattern of centrin (yellow). Scale bars are not adjusted to the expansion factor.

**Video S4:** Mitotic *Protodinium sp* stained against membranes using Bodipy (grey) and tubulin (magenta). Nuclear tunnels are shown, highlighting the presence of membranes around the tubulin spindle fibres. Scale bars are not adjusted to the expansion factor.

**Video S5:** *Heterocapsa rotundata* cell showing the apical centrin ring (yellow) and the cortical tubulin (magenta). Note that the ring is not fully closed and colocalises with tubulin below. Scale bars are not adjusted to the expansion factor.

**Video S6:** Apical region of a *Brandtodinium nutriculum* stained against centrin (yellow) and tubulin (magenta). Note the fully closed apical centrin ring that is partially colocalising with tubulin. Scale bars are not adjusted to the expansion factor.

**Video S7:** Centrin forms a cortical network in the cryptophyte *Rhodomonas sp*. Cell stained against centrin (yellow), tubulin (magenta) and DNA (blue). Scale bars are not adjusted to the expansion factor.

**Video S8:** Centrin forms the striated II fibres in *Pyramimonas diskoicola*. Tubulin (magenta) is staining the flagella and the cortical microtubules, while centrin (yellow) is decorating 2 main fibres transversally and is also present in the distal region of the basal bodies. Scale bars are not adjusted to the expansion factor.

**Video S9:** Structure of the oral apparatus of *Lacrymarya sp.* Cell was stained against centrin (yellow) and tubulin (magenta) and with NHS-ester staining (grey) and DNA (blue). Note the structure of centrin in a hexagonal shape in the body while in the oral apparatus forms a ring below the tornado shape tubulin fibres. Scale bars are not adjusted to the expansion factor.

**Video S10:** *Vorticella sp*. stained against tubulin (magenta), centrin (yellow) and DNA (blue). Note the structured organisation of both proteins in the oral apparatus. Scale bars are not adjusted to the expansion factor.

## Methods

### Cell chemical and cryofixation

All species cultures were obtained directly on site from the Roscoff Culture Collection (RCC) and the Basque Microalgae Culture Collection (BMCC). The cultures were maintained under their standard conditions as outlined on their respective websites (https://roscoff-culture-collection.org/ and https://www.ehu.eus/en/web/bmcc/search) respectively. For chemical fixation, cells were directly mixed with an equal volume of 8% formaldehyde (FA) solution, without any prior concentration step, to avoid cell stress and potential damage from centrifugation. This resulted in a final FA concentration of 4% (v/v). Fixation proceeded for 20 minutes at room temperature. Cells that did not settle during fixation were collected by centrifugation at 1,000g for 5 minutes, transferred to 1.5 mL tubes, and washed once with PBS. They were then collected again by centrifugation at 1,000g in a swing-out rotor and stored in 1% (v/v) FA in PBS at 4°C until further processing. In a few cases (Table S1), samples were vitrified using either high-pressure freezing (HPF) or plunge freezing. HPF was performed using a Leica EM ICE, where 1–1.5 µL of samples concentrated by brief centrifugation at 1,000g were loaded into hexadecene-coated 3 mm carriers. Multiple carriers were frozen for each sample to increase cell numbers and stored in liquid nitrogen. For plunge freezing, unconcentrated cells were loaded onto 12 mm coverslips coated with Poly-L-Lysine (Merck P4832) and CellTaK (Corning 354240) to promote cell adherence. After allowing the cells to settle for 10–15 minutes, excess liquid was removed, and any residual medium was blotted off using Whatman filter paper before rapidly plunging the coverslips into liquid ethane with a manual plunging system (RHOST LLC). Samples were stored in homemade metal coverslip racks in liquid nitrogen.

For both high-pressure and plunge-frozen samples, freeze substitution was required before expansion as previously described. ^26,27^ This was performed by placing the samples into frozen acetone containing 0.5% (v/v) FA and 0.025% (v/v) GA, followed by a gradual increase in temperature to room temperature. For HPF samples, an automated freeze substitution system (Leica AFS) was used, maintaining -90°C for at least 24 hours before increasing the temperature at a rate of 5°C per hour. For plunge-frozen samples, tubes containing coverslips in acetone were submerged in dry ice and incubated overnight. The dry ice was then incrementally removed over several hours to allow for a slow temperature increase. Subsequently, the samples were rehydrated by placing them in 100% ethanol, followed by a series of washes with ethanol in water (2x 95%, 1x 75%, 1x 50%, 3 minutes each). Finally, the samples were stored in PBS at 4°C until further use.^27^

### Ultrastructure Expansion Microscopy (U-ExM)

U-ExM was performed as previously described, with slight modifications.^15,21,25^ Formaldehyde-fixed cells (10 µL) were incubated in 500 µL anchoring solution (2% acrylamide, 1.4% formaldehyde) overnight at 37°C in PCR tubes. The following day, cells were placed onto either Poly-L-Lysine and CellTak or Poly-D-Lysine coated coverslips for 20–30 minutes at room temperature. Excess supernatant was removed, and the cells were embedded in a matrix gel composed of monomer solution (MS) [19% (wt/wt) sodium acrylate (Chem Cruz, Sigma 7446-81-3), 10% (wt/wt) acrylamide (Sigma-Aldrich A4058), and 0.1% (wt/wt) N,N’-methylenbisacrylamide (Sigma-Aldrich M1533) in PBS], with 0.5% (wt/v) TEMED and 0.5% (wt/v) APS added immediately before use. To slow down polymerization and allow sample placement and infiltration, those steps were performed on ice and the samples were incubated at 4°C for 5 minutes, then carefully transferred to a humidified box and incubated at 37°C for 45 minutes to 1 hour. After polymerization, gels were incubated in preheated denaturation buffer [50 mM Tris pH 9.0, 200 mM NaCl, 200 mM SDS, pH adjusted to 9.0 with HCl] for 1.5 hours at 95°C. Finally, gels were washed with water three times, and the size of the gel was measured to determine the expansion factor. For staining, expanded samples were shrunk by incubating them in PBS for 10 minutes, twice. In some cases, gels were cut into smaller pieces at this point to conserve material. For antibody staining, gels were incubated with primary antibodies (Table S2) in 3% BSA in PBS-T (0.2% (v/v) Tween-20 in PBS) for at least 4 hours to overnight at 37°C. Samples were then washed three times with PBS-T, 10 minutes per wash, before adding secondary antibodies (Table S3) in 3% (wt/v) BSA in PBS-T. Staining was performed at 37°C for at least 2.5 hours, followed by three washes with PBS-T for 10 minutes each. Protein pan-labelling with NHS ester [Dylight 405 (ThermoFisher, 46400), Alexa Fluor NHS-Ester 594 (ThermoFisher, A20004)] or membrane staining with Bodipy TR ceramide (ThermoFisher D7540, 2 mM stock in dimethyl sulfoxide, 1:2000 dilution) was performed together with DNA labelling using Hoechst 33342 (ThermoFisher 62249) or DAPI in PBS. Gels were washed once with PBS and then stained with the dyes diluted in PBS for 1–2 hours at 37°C or overnight at 4°C. During all incubations, tubes were kept on rotating wheels to ensure even staining. Once staining was completed, samples were washed twice with PBS for 5 minutes each. The samples were either expanded in H2O for imaging or stored in PBS at 4°C. For longer storage, gels were kept in PBS with 0.002% (wt/v) sodium azide at 4°C.For gel mounting, gels were cut to appropriate sizes and either attached to pre-coated PLL coverslips and sealed using i-Spacers (Sunjin Lab; #IS013) or placed onto PLL-coated ibidi glass bottom chambers. PLL will avoid drifting during acquisition.

**Table S2:** List of primary antibodies used in this study:

| Primary antibody | Source | Concentration |
| --- | --- | --- |
| *anti-tubulin, E7* | *DSHB* | 1:500 |
| *anti-tubulin, AA344 & AA345* | *ABCD antibodies* | 1:500 |
| *anti-NPC, Mab414* | *Biolegend 902901* | 1:250 |
| *Anti-centrin, 20H5* | *Millipore 04-1624* | 1:250 |
| *anti-histone 3* | *Agrisera AS10 710* | 1:500 |
| *GT335* | *Carsten Janke* | 1:500 |
| *Azo49* | *Carsten Janke* | 1:500 |
| *Azo49* | *Carsten Janke* | 1:500 |
| *Gly-Pep1* | *Carsten Janke* | 1:500 |

**Tables S3:**
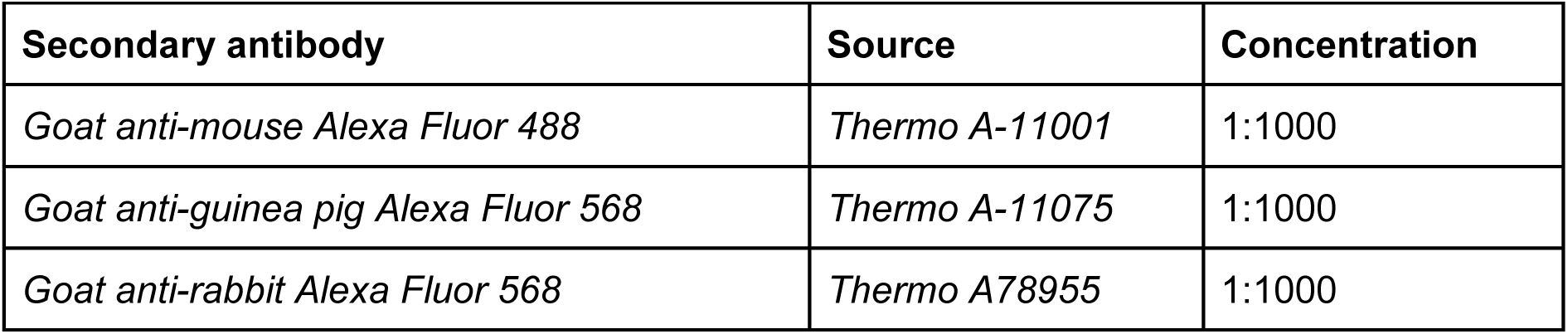
List of secondary antibodies used in this study:

| Secondary antibody | Source | Concentration |
| --- | --- | --- |
| Goat anti-mouse Alexa Fluor 488 | Thermo A-11001 | 1:1000 |
| Goat anti-guinea pig Alexa Fluor 568 | Thermo A-11075 | 1:1000 |
| Goat anti-rabbit Alexa Fluor 568 | Thermo A78955 | 1:1000 |

### Culturing of environmental samples and genotyping

Samples were collected in shallow water at 35° 37’ 53.1906’’, 139° 46’ 37.2072’ on 13.06.2023 and maintained in artificial sea water at 36 g/L with 0.1% enrichment medium (yeast extract 3 g/L, malt extract 3 g/L, peptone 5 g/L, glucose 10 g/L, NaCl 20 g/L). Flasks were kept at 23°C without shaking in the dark and split when required. *Euplotes rariseta* was isolated using serial dilutions and cell picking and maintained in artificial sea water with grains of autoclaved rice in the same conditions. Cell fixation was performed by adding 4% FA or 4% FA with 0.01% GA to the cells in medium and washing fixatives off after 20 min at RT. For amoeba, coverslips were placed into 6 well plates seeded from the cultures and grown over night. Coverslips were fixed by placing them into either 4% FA, ice cold methanol (for 5 min) or plunge frozen as described above. *Euplotes rariseta* and *Flabellula* sp. were identified by amplifying the 18S locus using oligos described and sending it for sequencing with GATC, other species according to their cellular morphology.^94^

### Microscopy

Image acquisition was performed using various microscopes. A Nikon-CSU-W1 SORA spinning disk confocal microscope was used with water immersion objectives (Apo LWD 40x WI Lambda-S/1.15, 0.60 and SR P-Apochromat IR AC 60x WI/1.27) and a CSUW1-TS 2D 50/SoRa disk. In some cases, an additional 4x SoRa disk was employed to further increase magnification. Alternatively, an Olympus IXplore SpinSR spinning disk microscope, also equipped with a CSUW1-TS 2D 50/SoRa disk, was used with a UPLSAPO 10x/NA 0.4 air objective for overviews and a PLAPON 60x/NA 1.42 oil objective for high-resolution imaging. For AiryScan microscopy, samples were imaged on a Zeiss LSM980 Airy Fast with a Plan-Apochromat 63x/1.4 Oil DIC M27 objective. Imaging was also performed on a Nikon A1R HD25 laser scanning confocal microscope with CFI Apo LWD Lambda-S 40x/1.15 WI, 0.60 and CFI Plan Apochromat VC 60x/1.3 WI objectives. Additionally, a Leica Stellaris 8 Falcon microscope with a 63x/1.4 NA oil immersion objective and an upright Leica SP8 confocal microscope with an HC PL APO 40x/1.25 glycerol objective were used. Widefield imaging was conducted on a Leica Thunder DMi8 microscope with a 63x/1.4 NA oil immersion objective, and the images were deconvoluted using Leica Thunder SVCC software with ‘Water’ as the mounting medium. For STED microscopy, a Leica SP8 STED 3X equipped with a PL Apo CS2 100x Oil NA 1.4 objective was used, while a Nikon AX-NSPARC with Apo LWD 40x WI NA 1.15 and PLAN Apo VC 60xA WI NA 1.2 objectives were used for few samples.

### Image analysis

Images were processed and analysed using FIJI. ^34,35^ The foundational script for line plot analysis was generated with the assistance of the Large Language Model ChatGPT-4 (OpenAI, version 2 September 2024) and later finalized and analysed in RStudio (RStudio Team (2020). RStudio: Integrated Development for R. RStudio, PBC, Boston, MA URL http://www.rstudio.com/). Excel was used to determine the distance between peaks. For analysis of MT array (Figure 2B and 2C) and chromosome band spacing (Figure 3B) in dinoflagellates, the PatternJ toolset was used. Briefly, a line was drawn across a region of the nucleus or cell to extract peaks positions and analyse spacing between adjacent peaks. The values obtained were adjusted for expansion factor. Graphs were prepared in GraphPad Prism 9 or 10.3.1 (GraphPad Software, Boston, Massachusetts USA, www.graphpad.com). All the 3D reconstructions were made in Imaris 10.1.0 (Oxford Instruments). In all cases, the scale bars are not adjusted to the expansion factor. Segmentation and visualisation of volumes of *Urgorri complanatus* (Figure 1) were performed in Amira (v.2019.3 or 2020.1; ThermoFisher Scientific). For the analysis of hexagon areas in *Rhinomonas nottbecki*, hexagons were manually drawn by connecting centrin dots. A mask was then applied to calculate the area of each hexagon. For the analysis of the apical ring in dinoflagellates, two different approaches were used. The first approach involved using FIJI to analyze the diameter of the lateral view of centrin rings. For measuring the perimeter, 3D reconstructions of each ring were created using Imaris 10.1.0, after which the perimeter was calculated.

### Resource sharing using MoBIE

To make the images easily and openly accessible, we relied on the ImageJ - plugin MoBIE. ^36^ A MoBIE project was created by first converting image volumes to the ome-zarr file format using the bioformats2raw (https://github.com/glencoesoftware/bioformats2raw) command line tool with a chunk size of 96 x 96 x 96. The converted images were then added to the MoBIE project using the MoBIE python API (https://github.com/mobie/mobie-utils-python/tree/master) and uploaded to an S3 bucket hosted by EMBL (https://console.s3.embl.de/browser/culture-collections) via the MinIO client (https://min.io/docs/minio/linux/reference/minio-mc.html). All steps were automated using Python scripts, which are available along with the MoBIE project metadata in the following GitHUB repository: https://github.com/Dey-Lab-Team/culture-collections. The data can be downloaded from the S3 bucket (https://console.s3.embl.de/browser/culture-collections) or visualised directly in MoBIE. To do this, install Fiji (https://imagej.net/software/fiji/downloads) and the MoBIE plugin (https://mobie.github.io/tutorials/installation.html). Start MoBIE by searching for “mobie” in the search bar and running the “Open MoBIE Project…” workflow. Then, enter the GitHub repository link (https://console.s3.embl.de/browser/culture-collections) and choose “Remote”.

## References

1. Lane, N. (2015). The unseen world: reflections on Leeuwenhoek (1677) ‘Concerning little animals.’ Phil. Trans. R. Soc. B 370, 20140344. 10.1098/rstb.2014.0344.

2. Deacon, M., Rice, A.L., and Summerhayes, C.P. (2013). Understanding the oceans: a century of ocean exploration (Routledge).

3. Aitken, F., Foulc, J.-N., and Aitken, F. (2019). The first explorations on the deep sea by H.M.S. Challenger (1872-1876) (ISTE Ltd).

4. Levit, G.S., and Hossfeld, U. (2019). Ernst Haeckel in the history of biology. Current Biology 29, R1276–R1284. 10.1016/j.cub.2019.10.064.

5. Dobell, C. (1923). A Protozoological Bicentenary: Antony van Leeuwenhoek (1632–1723) and Louis Joblot (1645–1723). Parasitology 15, 308–319. 10.1017/S0031182000014797.

6. Burki, F., Sandin, M.M., and Jamy, M. (2021). Diversity and ecology of protists revealed by metabarcoding. Current Biology 31, R1267–R1280. 10.1016/j.cub.2021.07.066.

7. Bjorbækmo, M.F.M., Evenstad, A., Røsæg, L.L., Krabberød, A.K., and Logares, R. (2020). The planktonic protist interactome: where do we stand after a century of research? The ISME Journal 14, 544–559. 10.1038/s41396-019-0542-5.

8. New, F.N., and Brito, I.L. (2020). What Is Metagenomics Teaching Us, and What Is Missed? Annu. Rev. Microbiol. 74, 117–135. 10.1146/annurev-micro-012520-072314.

9. Colin, S., Coelho, L.P., Sunagawa, S., Bowler, C., Karsenti, E., Bork, P., Pepperkok, R., and De Vargas, C. (2017). Quantitative 3D-imaging for cell biology and ecology of environmental microbial eukaryotes. eLife 6, e26066. 10.7554/eLife.26066.

10. Hirst, M.B., Kita, K.N., and Dawson, S.C. (2011). Uncultivated Microbial Eukaryotic Diversity: A Method to Link ssu rRNA Gene Sequences with Morphology. PLoS ONE 6, e28158. 10.1371/journal.pone.0028158.

11. Collinson, L.M., Bosch, C., Bullen, A., Burden, J.J., Carzaniga, R., Cheng, C., Darrow, M.C., Fletcher, G., Johnson, E., Narayan, K., et al. (2023). Volume EM: a quiet revolution takes shape. Nat Methods 20, 777–782. 10.1038/s41592-023-01861-8.

12. Chen, F., Tillberg, P.W., and Boyden, E.S. (2015). Expansion microscopy. Science 347, 543–548. 10.1126/science.1260088.

13. Chozinski, T.J., Halpern, A.R., Okawa, H., Kim, H.-J., Tremel, G.J., Wong, R.O.L., and Vaughan, J.C. (2016). Expansion microscopy with conventional antibodies and fluorescent proteins. Nat Methods 13, 485–488. 10.1038/nmeth.3833.

14. Tillberg, P.W., Chen, F., Piatkevich, K.D., Zhao, Y., Yu, C.-C., English, B.P., Gao, L., Martorell, A., Suk, H.-J., Yoshida, F., et al. (2016). Protein-retention expansion microscopy of cells and tissues labeled using standard fluorescent proteins and antibodies. Nat Biotechnol 34, 987–992. 10.1038/nbt.3625.

15. Gambarotto, D., Zwettler, F.U., Le Guennec, M., Schmidt-Cernohorska, M., Fortun, D., Borgers, S., Heine, J., Schloetel, J.-G., Reuss, M., Unser, M., et al. (2019). Imaging cellular ultrastructures using expansion microscopy (U-ExM). Nat Methods 16, 71–74. 10.1038/s41592-018-0238-1.

16. Cahoon, C.K., Yu, Z., Wang, Y., Guo, F., Unruh, J.R., Slaughter, B.D., and Hawley, R.S. (2017). Superresolution expansion microscopy reveals the three-dimensional organization of the *Drosophila* synaptonemal complex. Proc. Natl. Acad. Sci. U.S.A. 114. 10.1073/pnas.1705623114.

17. Freifeld, L., Odstrcil, I., Förster, D., Ramirez, A., Gagnon, J.A., Randlett, O., Costa, E.K., Asano, S., Celiker, O.T., Gao, R., et al. (2017). Expansion microscopy of zebrafish for neuroscience and developmental biology studies. Proc. Natl. Acad. Sci. U.S.A. 114. 10.1073/pnas.1706281114.

18. Yu, C.-C. (Jay), Barry, N.C., Wassie, A.T., Sinha, A., Bhattacharya, A., Asano, S., Zhang, C., Chen, F., Hobert, O., Goodman, M.B., et al. (2020). Expansion microscopy of C. elegans. eLife 9, e46249. 10.7554/eLife.46249.

19. Bos, P.R., Berentsen, J., and Wientjes, E. (2023). Expansion microscopy resolves the thylakoid structure of spinach. Plant Physiology 194, 347–358. 10.1093/plphys/kiad526.

20. Laporte, M.H., Gambarotto, D., Bertiaux, É., Bournonville, L., Louvel, V., Nunes, J.M., Borgers, S., Hamel, V., and Guichard, P. (2024). Time-series reconstruction of the molecular architecture of human centriole assembly. Cell 187, 2158–2174.e19. 10.1016/j.cell.2024.03.025.

21. Tosetti, N., Dos Santos Pacheco, N., Bertiaux, E., Maco, B., Bournonville, L., Hamel, V., Guichard, P., and Soldati-Favre, D. (2020). Essential function of the alveolin network in the subpellicular microtubules and conoid assembly in Toxoplasma gondii. eLife 9, e56635. 10.7554/eLife.56635.

22. Bertiaux, E., Balestra, A.C., Bournonville, L., Louvel, V., Maco, B., Soldati-Favre, D., Brochet, M., Guichard, P., and Hamel, V. (2021). Expansion microscopy provides new insights into the cytoskeleton of malaria parasites including the conservation of a conoid. PLoS Biol 19, e3001020. 10.1371/journal.pbio.3001020.

23. Liffner, B., Cepeda Diaz, A.K., Blauwkamp, J., Anaguano, D., Frolich, S., Muralidharan, V., Wilson, D.W., Dvorin, J.D., and Absalon, S. (2023). Atlas of Plasmodium falciparum intraerythrocytic development using expansion microscopy. Elife 12, RP88088. 10.7554/eLife.88088.

24. Li, J., Shami, G.J., Liffner, B., Cho, E., Braet, F., Duraisingh, M.T., Absalon, S., Dixon, M.W.A., and Tilley, L. (2024). Disruption of Plasmodium falciparum kinetochore proteins destabilises the nexus between the centrosome equivalent and the mitotic apparatus. Nat Commun 15, 5794. 10.1038/s41467-024-50167-6.

25. Klena, N., Maltinti, G., Batman, U., Pigino, G., Guichard, P., and Hamel, V. (2023). An In-depth Guide to the Ultrastructural Expansion Microscopy (U-ExM) of Chlamydomonas reinhardtii. BIO-PROTOCOL 13. 10.21769/BioProtoc.4792.

26. Hinterndorfer, K., Laporte, M.H., Mikus, F., Tafur, L., Bourgoint, C., Prouteau, M., Dey, G., Loewith, R., Guichard, P., and Hamel, V. (2022). Ultrastructure expansion microscopy reveals the cellular architecture of budding and fission yeast. Journal of Cell Science 135, jcs260240. 10.1242/jcs.260240.

27. Laporte, M.H., Klena, N., Hamel, V., and Guichard, P. (2022). Visualizing the native cellular organization by coupling cryofixation with expansion microscopy (Cryo-ExM). Nat Methods 19, 216–222. 10.1038/s41592-021-01356-4.

28. Shah, H., Olivetta, M., Bhickta, C., Ronchi, P., Trupinić, M., Tromer, E.C., Tolić, I.M., Schwab, Y., Dudin, O., and Dey, G. (2024). Life-cycle-coupled evolution of mitosis in close relatives of animals. Nature 630, 116–122. 10.1038/s41586-024-07430-z.

29. Reza, M.H., Dutta, S., Goyal, R., Shah, H., Dey, G., and Sanyal, K. (2024). Expansion microscopy reveals characteristic ultrastructural features of pathogenic budding yeast species. J Cell Sci, jcs.262046. 10.1242/jcs.262046.

30. Bondarenko, K., Limoge, F., Pedram, K., Gissot, M., and Young, J.C. (2024). Enzymatically enhanced ultrastructure expansion microscopy unlocks expansion of in vitro Toxoplasma gondii cysts. mSphere 9, e0032224. 10.1128/msphere.00322-24.

31. Olivetta, M., Bhickta, C., Chiaruttini, N., Burns, J., and Dudin, O. (2024). A multicellular developmental program in a close animal relative. Preprint at Developmental Biology, 10.1101/2024.03.25.586530 https://doi.org/10.1101/2024.03.25.586530.

32. Vaulot, D., Gall, F., Le Marie, D., Guillou, L., and Partensky, F. (2004). The Roscoff Culture Collection (RCC): a collection dedicated to marine picoplankton. nova_hedwigia 79, 49–70. 10.1127/0029-5035/2004/0079-0049.

33. M’Saad, O., and Bewersdorf, J. (2020). Light microscopy of proteins in their ultrastructural context. Nat Commun 11, 3850. 10.1038/s41467-020-17523-8.

34. Schindelin, J., Arganda-Carreras, I., Frise, E., Kaynig, V., Longair, M., Pietzsch, T., Preibisch, S., Rueden, C., Saalfeld, S., Schmid, B., et al. (2012). Fiji: an open-source platform for biological-image analysis. Nat Methods 9, 676–682. 10.1038/nmeth.2019.

35. Rueden, C.T., Schindelin, J., Hiner, M.C., DeZonia, B.E., Walter, A.E., Arena, E.T., and Eliceiri, K.W. (2017). ImageJ2: ImageJ for the next generation of scientific image data. BMC Bioinformatics 18, 529. 10.1186/s12859-017-1934-z.

36. Pape, C., Meechan, K., Moreva, E., Schorb, M., Chiaruttini, N., Zinchenko, V., Martinez Vergara, H., Mizzon, G., Moore, J., Arendt, D., et al. (2023). MoBIE: a Fiji plugin for sharing and exploration of multi-modal cloud-hosted big image data. Nat Methods 20, 475–476. 10.1038/s41592-023-01776-4.

37. Gu, H., Luo, Z., Krock, B., Witt, M., and Tillmann, U. (2013). Morphology, phylogeny and azaspiracid profile of Azadinium poporum (Dinophyceae) from the China Sea. Harmful Algae 21–22, 64–75. 10.1016/j.hal.2012.11.009.

38. Mocaer, K., Mizzon, G., Gunkel, M., Halavatyi, A., Steyer, A., Oorschot, V., Schorb, M., Le Kieffre, C., Yee, D.P., Chevalier, F., et al. (2023). Targeted volume correlative light and electron microscopy of an environmental marine microorganism. J Cell Sci 136, jcs261355. 10.1242/jcs.261355.

39. Fenchel, T. (2001). How Dinoflagellates Swim. Protist 152, 329–338. 10.1078/1434-4610-00071.

40. Roberts, K.R., Lemoine, J.E., Schneider, R.M., and Farmer, M.A. (1988). The microtubular cytoskeleton of three dinoflagellates: an immunofluorescence study. Protoplasma 144, 68–71. 10.1007/BF01320283.

41. De Salas, M.F., Bolch, C.J.S., Botes, L., Nash, G., Wright, S.W., and Hallegraeff, G.M. (2003). *TAKAYAMA* GEN. NOV. (GYMNODINIALES, DINOPHYCEAE), A NEW GENUS OF UNARMORED DINOFLAGELLATES WITH SIGMOID APICAL GROOVES, INCLUDING THE DESCRIPTION OF TWO NEW SPECIES ^1^. Journal of Phycology 39, 1233–1246. 10.1111/j.0022-3646.2003.03-019.x.

42. Okamoto, N., and Keeling, P. (2014). A Comparative Overview of the Flagellar Apparatus of Dinoflagellate, Perkinsids and Colpodellids. Microorganisms 2, 73–91. 10.3390/microorganisms2010073.

43. Kusel-Fetzmann, E., and Weidinger, M. (2008). Ultrastructure of five Euglena species positioned in the subdivision Serpentes. Protoplasma 233, 209–222. 10.1007/s00709-008-0005-8.

44. Tashyreva, D., Týč, J., Horák, A., and Lukeš, J. (2023). Ultrastructure and 3D reconstruction of a diplonemid protist (Diplonemea) and its novel membranous organelle. mBio 14, e01921–23. 10.1128/mbio.01921-23.

45. Perret, E., Albert, M., Bordes, N., Bornens, M., and Soyer-Gobillard, M.-O. (1991). Microtubular spindle and centrosome structures during the cell cycle in a dinoflagellate Crypthecodinium cohnii B.: an immunocytochemical study. Biosystems 25, 53–65. 10.1016/0303-2647(91)90012-A.

46. Velasquez-Carvajal, D., Garampon, F., Besnardeau, L., Lemée, R., Schaub, S., and Castagnetti, S. (2024). Microtubule reorganization during mitotic cell division in the dinoflagellate *Ostreospis* cf. *ovata*. Journal of Cell Science 137, jcs261733. 10.1242/jcs.261733.

47. Bré, M.-H., Redeker, V., Quibell, M., Darmanaden-Delorme, J., Bressac, C., Cosson, J., Huitorel, P., Schmitter, J.-M., Rossier, J., Johnson, T., et al. (1996). Axonemal tubulin polyglycylation probed with two monoclonal antibodies: widespread evolutionary distribution, appearance during spermatozoan maturation and possible function in motility. Journal of Cell Science 109, 727–738. 10.1242/jcs.109.4.727.

48. Janouškovec, J., Gavelis, G.S., Burki, F., Dinh, D., Bachvaroff, T.R., Gornik, S.G., Bright, K.J., Imanian, B., Strom, S.L., Delwiche, C.F., et al. (2017). Major transitions in dinoflagellate evolution unveiled by phylotranscriptomics. Proc. Natl. Acad. Sci. U.S.A. 114. 10.1073/pnas.1614842114.

49. Taylor, F.J.R. (1980). On dinoflagellate evolution. Biosystems 13, 65–108. 10.1016/0303-2647(80)90006-4.

50. Gauthier, G.M. (2015). Dimorphism in Fungal Pathogens of Mammals, Plants, and Insects. PLoS Pathog 11, e1004608. 10.1371/journal.ppat.1004608.

51. Gornik, S.G., Ford, K.L., Mulhern, T.D., Bacic, A., McFadden, G.I., and Waller, R.F. (2012). Loss of Nucleosomal DNA Condensation Coincides with Appearance of a Novel Nuclear Protein in Dinoflagellates. Current Biology 22, 2303–2312. 10.1016/j.cub.2012.10.036.

52. Irwin, N.A.T., Martin, B.J.E., Young, B.P., Browne, M.J.G., Flaus, A., Loewen, C.J.R., Keeling, P.J., and Howe, L.J. (2018). Viral proteins as a potential driver of histone depletion in dinoflagellates. Nat Commun 9, 1535. 10.1038/s41467-018-03993-4.

53. Gavelis, G.S., Herranz, M., Wakeman, K.C., Ripken, C., Mitarai, S., Gile, G.H., Keeling, P.J., and Leander, B.S. (2019). Dinoflagellate nucleus contains an extensive endomembrane network, the nuclear net. Sci Rep 9, 839. 10.1038/s41598-018-37065-w.

54. Drechsler, H., and McAinsh, A.D. (2012). Exotic mitotic mechanisms. Open Biol. 2, 120140. 10.1098/rsob.120140.

55. Salisbury, J.L., Baron, A., Surek, B., and Melkonian, M. (1984). Striated flagellar roots: isolation and partial characterization of a calcium-modulated contractile organelle. The Journal of cell biology 99, 962–970. 10.1083/jcb.99.3.962.

56. Melkonian, M., Beech, P.L., Katsaros, C., and Schulze, D. (1992). Centrin-Mediated Cell Motility in Algae. In Algal Cell Motility, M. Melkonian, ed. (Springer US), pp. 179–221. 10.1007/978-1-4615-9683-7_6.

57. Salisbury, J.L. (1995). Centrin, centrosomes, and mitotic spindle poles. Current Opinion in Cell Biology 7, 39–45. 10.1016/0955-0674(95)80043-3.

58. Levy, Y.Y., Lai, E.Y., Remillard, S.P., Heintzelman, M.B., and Fulton, C. (1996). Centrin is a conserved protein that forms diverse associations with centrioles and MTOCs inNaegleria and other organisms. Cell Motil. Cytoskeleton 33, 298–323. 10.1002/(SICI)1097-0169(1996)33:4<298::AID-CM6>3.0.CO;2-5.

59. Schiebel, E., and Bornens, M. (1995). In search of a function for centrins. Trends in Cell Biology 5, 197–201. 10.1016/0962-8924(95)80011-5.

60. Huang, B., Watterson, D.M., Lee, V.D., and Schibler, M.J. (1988). Purification and characterization of a basal body-associated Ca2+-binding protein. J Cell Biol 107, 121–131. 10.1083/jcb.107.1.121.

61. Tourbez, M., Firanescu, C., Yang, A., Unipan, L., Duchambon, P., Blouquit, Y., and Craescu, C.T. (2004). Calcium-dependent Self-assembly of Human Centrin 2. Journal of Biological Chemistry 279, 47672–47680. 10.1074/jbc.M404996200.

62. Diogon, M., Henou, C., Ravet, V., Bouchard, P., and Viguès, B. (2001). Evidence for regional differences in the dynamicsof centrin cytoskeletal structures in the polymorphichymenostome ciliate Tetrahymena paravorax. European Journal of Protistology 37, 223–231. 10.1078/0932-4739-00824.

63. Dantas, T.J., Daly, O.M., and Morrison, C.G. (2012). Such small hands: the roles of centrins/caltractins in the centriole and in genome maintenance. Cell. Mol. Life Sci. 69, 2979–2997. 10.1007/s00018-012-0961-1.

64. Schweizer, N., Haren, L., Dutto, I., Viais, R., Lacasa, C., Merdes, A., and Lüders, J. (2021). Sub-centrosomal mapping identifies augmin-γTuRC as part of a centriole-stabilizing scaffold. Nat Commun 12, 6042. 10.1038/s41467-021-26252-5.

65. Kalichava, A., and Ochsenreiter, T. (2021). Ultrastructure Expansion Microscopy in *Trypanosoma brucei* (Cell Biology) 10.1101/2021.04.20.440568.

66. Höhfeld, I., Otten, J., and Melkonian, M. (1988). Contractile eukaryotic flagella: Centrin is involved. Protoplasma 147, 16–24. 10.1007/BF01403874.

67. Maruyama, T. (1982). Fine structure of the longitudinal flagellum in *Ceratium tripos* , a marine dinoflagellate. Journal of Cell Science 58, 109–123. 10.1242/jcs.58.1.109.

68. Roberts, K.R., Timpano, P., and Montegut, A.E. (1987). The apical pore fibrous complex: a new cytological feature of some dinoflagellates. Protoplasma 137, 65–69. 10.1007/BF01281178.

69. De Souza, W., and Attias, M. (2010). Subpellicular Microtubules in Apicomplexa and Trypanosomatids. In Structures and Organelles in Pathogenic Protists Microbiology Monographs., W. De Souza, ed. (Springer Berlin Heidelberg), pp. 27–62. 10.1007/978-3-642-12863-9_2.

70. Dos Santos Pacheco, N., Tosetti, N., Koreny, L., Waller, R.F., and Soldati-Favre, D. (2020). Evolution, Composition, Assembly, and Function of the Conoid in Apicomplexa. Trends in Parasitology 36, 688–704. 10.1016/j.pt.2020.05.001.

71. Koreny, L., Zeeshan, M., Barylyuk, K., Tromer, E.C., Van Hooff, J.J.E., Brady, D., Ke, H., Chelaghma, S., Ferguson, D.J.P., Eme, L., et al. (2021). Molecular characterization of the conoid complex in Toxoplasma reveals its conservation in all apicomplexans, including Plasmodium species. PLoS Biol 19, e3001081. 10.1371/journal.pbio.3001081.

72. Waller, R.F., and Carruthers, V.B. (2024). Adaptations and metabolic evolution of myzozoan protists across diverse lifestyles and environments. Microbiol Mol Biol Rev, e00197–22. 10.1128/mmbr.00197-22.

73. Leander, B.S., and Keeling, P.J. (2003). Morphostasis in alveolate evolution. Trends in Ecology & Evolution 18, 395–402. 10.1016/S0169-5347(03)00152-6.

74. Gubbels, M.-J., and Duraisingh, M.T. (2012). Evolution of apicomplexan secretory organelles. International Journal for Parasitology 42, 1071–1081. 10.1016/j.ijpara.2012.09.009.

75. Geimer, S., and Melkonian, M. (2005). Centrin Scaffold in *Chlamydomonas reinhardtii* Revealed by Immunoelectron Microscopy. Eukaryot Cell 4, 1253–1263. 10.1128/EC.4.7.1253-1263.2005.

76. Hill, D.R.A., and Wetherbee, R. (1988). The structure and taxonomy of Rhinomonas pauca gen. et sp. nov. (Cryptophyceae). Phycologia 27, 355–365. 10.2216/i0031-8884-27-3-355.1.

77. Salisbury, J.L., Swanson, J.A., Floyd, G.L., Hall, R., and Maihle, N.J. (1981). Ultrastructure of the flagellar apparatus of the green algaTetraselmis subcordiformis: With special consideration given to the function of the rhizoplast and rhizanchora. Protoplasma 107, 1–11. 10.1007/BF01275602.

78. Lechtreck, K.-F., and Melkonian, M. (1991). An update on fibrous flagellar roots in green algae. Protoplasma 164, 38–44. 10.1007/BF01320813.

79. Allen, R.L., Kennel, S.J., Cacheiro, L., Olins, A.L., and Olins, D.E. (1986). Examination of the macronuclear replication band in Euplotes eurystomus with monoclonal antibodies. The Journal of cell biology 102, 131–136. 10.1083/jcb.102.1.131.

80. Fu, J., Chi, Y., Lu, X., Gao, F., Al-Farraj, S.A., Petroni, G., and Jiang, J. (2022). Doublets of the unicellular organism Euplotes vannus (Alveolata, Ciliophora, Euplotida): the morphogenetic patterns of the ciliary and nuclear apparatuses associated with cell division. Mar Life Sci Technol 4, 527–535. 10.1007/s42995-022-00150-1.

81. Coyle, S.M., Flaum, E.M., Li, H., Krishnamurthy, D., and Prakash, M. (2019). Coupled Active Systems Encode an Emergent Hunting Behavior in the Unicellular Predator Lacrymaria olor. Current Biology 29, 3838–3850.e3. 10.1016/j.cub.2019.09.034.

82. Qin, W., Hu, C., Gu, S., Zhang, J., Jiang, C., Chai, X., Liao, Z., Yang, M., Zhou, F., Kang, D., et al. (2024). Dynamic shape-shifting of the single-celled eukaryotic predator Lacrymaria via unconventional cytoskeletal components. Current Biology, S0960982224012144. 10.1016/j.cub.2024.09.003.

83. Allen, R. (2021). CIL:39510, Vorticella convallaria, cell by organism, eukaryotic cell, Eukaryotic Protist, Ciliated Protist. In Cell Image Library. (UC San Diego Library Digital Collections). 10.6075/J0930S2P https://doi.org/10.6075/J0930S2P.

84. Amos, W.B. (1972). Structure and coiling of the stalk in the peritrich ciliates *Vorticella* and *Carchesium*. Journal of Cell Science 10, 95–122. 10.1242/jcs.10.1.95.

85. Allen, R.D. (1973). STRUCTURES LINKING THE MYONEMES, ENDOPLASMIC RETICULUM, AND SURFACE MEMBRANES IN THE CONTRACTILE CILIATE *VORTICELLA*. The Journal of Cell Biology 56, 559–579. 10.1083/jcb.56.2.559.

86. Wall, R.J., Roques, M., Katris, N.J., Koreny, L., Stanway, R.R., Brady, D., Waller, R.F., and Tewari, R. (2016). SAS6-like protein in Plasmodium indicates that conoid-associated apical complex proteins persist in invasive stages within the mosquito vector. Sci Rep 6, 28604. 10.1038/srep28604.

87. Vonderfecht, T., Stemm-Wolf, A.J., Hendershott, M., Giddings, T.H., Meehl, J.B., and Winey, M. (2011). The two domains of centrin have distinct basal body functions in Tetrahymena. Mol Biol Cell 22, 2221–2234. 10.1091/mbc.E11-02-0151.

88. Guerra, C., Wada, Y., Leick, V., Bell, A., and Satir, P. (2003). Cloning, localization, and axonemal function of Tetrahymena centrin. Mol Biol Cell 14, 251–261. 10.1091/mbc.e02-05-0298.

89. Pedretti, M., Bombardi, L., Conter, C., Favretto, F., Dominici, P., and Astegno, A. (2021). Structural Basis for the Functional Diversity of Centrins: A Focus on Calcium Sensing Properties and Target Recognition. Int J Mol Sci 22, 12173. 10.3390/ijms222212173.

90. Dawson, S.C., and Fritz-Laylin, L.K. (2009). Sequencing free-living protists: the case for metagenomics. Environ Microbiol 11, 1627–1631. 10.1111/j.1462-2920.2009.01965.x.

91. Griffiths, G. (1993). Fine Structure Immunocytochemistry (Springer Berlin Heidelberg) 10.1007/978-3-642-77095-1.

92. Cooney, E.C., Holt, C.C., Hehenberger, E., Adams, J.A., Leander, B.S., and Keeling, P.J. (2024). Investigation of heterotrophs reveals new insights in dinoflagellate evolution. Molecular Phylogenetics and Evolution 196, 108086. 10.1016/j.ympev.2024.108086.

93. Novák Vanclová, A.M., Nef, C., Füssy, Z., Vancl, A., Liu, F., Bowler, C., and Dorrell, R.G. (2024). New plastids, old proteins: repeated endosymbiotic acquisitions in kareniacean dinoflagellates. EMBO Rep 25, 1859–1885. 10.1038/s44319-024-00103-y.

94. Šlapeta, J., Moreira, D., and López-García, P. (2005). The extent of protist diversity: insights from molecular ecology of freshwater eukaryotes. Proc. R. Soc. B. 272, 2073– 2081. 10.1098/rspb.2005.3195.

